# Accurate Identification of Functional Residues Across the Human Proteome with TAMALE

**DOI:** 10.64898/2026.05.28.728550

**Authors:** Justin Van Riper, Bridget J. Corsaro, Monica C. Pillon

## Abstract

The central challenge of the post-AlphaFold era is the ‘functional gap’. Despite having structure predictions for nearly every human protein, we remain unable to systematically distinguish residues that drive activity from those that merely maintain structural integrity. Here we introduce TAMALE, a machine learning model that calculates graded residue-level functional scores across the human proteome without prior annotation. TAMALE transforms structure models and variant effect predictions to identify residues involved in catalysis, ligand binding, nucleic acid interactions, and regulation. The model also distinguishes pseudo-enzymes from catalytically active homologs. The model was validated across 20 case studies along with experimental characterization of the FASTKD5 ribonuclease, demonstrating its utility for functional discovery. Applied proteome-wide to 19,528 human proteins, TAMALE generates testable hypotheses enabling mechanistic discovery at scale.

## Main Text

Proteins underpin the molecular basis of biology through spatially organized functional sites that mediate chemical reactions, molecular interactions, and regulation. A major challenge of the post-AlphaFold era is the inability to systematically distinguish residues that drive activity from those that maintain structural integrity (*1*). Incomplete annotation, lineage-specific adaptations, and technical limitations leave the human proteome disproportionately unresolved, restricting mechanistic insight into fundamental biology and therapeutics.

Computational strategies for predicting functional residues remain limited. Sequence- and structure-based approaches infer function from characterized homologs but assume similarity implies functional equivalence (*2, 3*). Hybrid approaches integrate multiple predefined features but obscure the distinction of functional clusters from structural constraints (*4, 5*). More recently, machine-learning models have improved residue-level prediction by learning context-dependent patterns (*6-8*). Notably, EasiFA incorporates catalytic reaction information for enzyme annotation, but its reliance on known mechanisms limits its ability to detect novel biology and reduces its capacity to identify non-catalytic functional sites (*6*).

AlphaMissense (Missense) provides proteome-wide pathogenicity predictions for all possible amino acid substitutions, offering an underexploited resource for functional residue identification (*9*). Missense integrates signals derived from allele frequency, conservation, co-evolution, and structural context (*1, 9*). Although Missense is not explicitly trained to identify residues with functional roles, we hypothesize that it contains latent biochemical constraints that can be decoded into residue-level functional information across the human proteome.

In this study, we present TAMALE (<u>T</u>argeted <u>A</u>lpha<u>M</u>issense <u>A</u>nalysis in <u>L</u>ocal structural <u>E</u>nvironments) which leverages local structural context to transform variant effect predictions into graded residue-level functionality scores. This machine-learning model selectively predicts functionally important residues over positions whose sensitivity primarily arises from structural roles. TAMALE enriches for residues involved in catalysis, metal ion coordination, ligand binding, peptide and nucleic acid interactions, and regulatory or allosteric positions. The model was benchmarked on 453 enzymes, and we present 20 case studies of previously characterized proteins along with an experimental discovery case of the FASTKD5 ribonuclease. TAMALE does not require prior functional annotation, enabling a scalable framework for systematic discovery of catalytic and non-catalytic functional sites across the human proteome.

### TAMALE architecture

TAMALE integrates structure predictions, variant effect scores, and physicochemical properties to infer residue-level functional importance. Using the UniProt identifier as the input, the algorithm retrieves the structure prediction, associated predicted aligned error (PAE) estimates, and variant effect pathogenicity scores from the AlphaFold database (**Fig. 1A**) (*1, 9, 10*). Forty-eight residue-level features are derived to integrate structural geometry, evolutionary constraints, and variant effect signals (**table S1**, see materials and methods). TAMALE repurposes Missense pathogenicity predictions, typically used for disease interpretation, to capture latent biochemical constraints associated with the sensitivity and spatial clustering of each amino acid. Together, these inputs enable systematic prioritization of functional residues over structural positions.

**Fig. 1.**
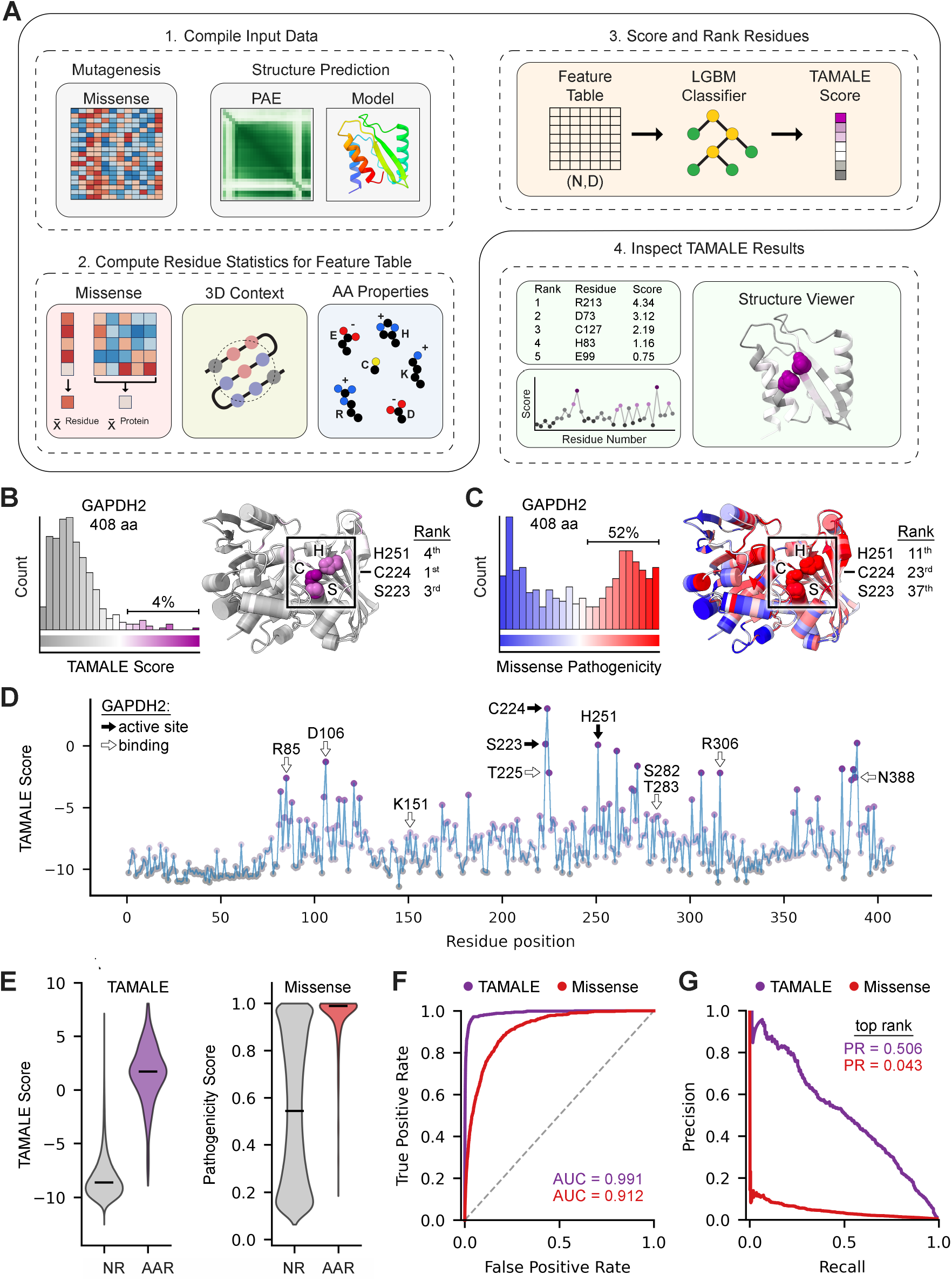
TAMALE exhibits robust active site selectivity. (**A**) Schematic of TAMALE for predicting direct functional residues. (**B**) Prediction of GAPDH2 active site residues based on TAMALE scores. Graph quantifies the distribution of ranked residues across the score range and the AlphaFold (AF) model illustrates the distribution of scores, where purple predicts functionally important residues and grey is likely nonfunctional residues. Annotated active site residues are boxed and listed with their respective ranked position. (**C**) Same as panel B but using mean pathogenicity where red are sensitive residue positions (pathogenic) and blue are insensitive positions (benign) (*1, 9, 12*). (**D**) GAPDH2 sequence track of TAMALE scores with annotated active site (black arrow) and binding residues (white arrow) are marked for reference (*10*). (**E**) Violin plots depict the score distribution between annotated active site residues (AAR) and non-annotated residues (NR) based on TAMALE score and mean pathogenicity. (**F**) Receiver operating characteristic curve with an area under the curve of 0.991 for TAMALE and 0.912 for Missense. Dotted line is random chance. (**G**) Precision-recall curves comparing the performance of TAMALE and Missense in predicting an annotated active site residue in its top ranked position.

TAMALE was trained to predict functionally important residues using 1,356 reviewed human enzymes representing all general enzyme commission (EC) numbers (**data S1**) (*10*). Functional labels were derived from active site annotations listed in the UniProt database, including catalytic and ligand binding residues (*10*). In total, 2,142 annotated active site residues served as proxy functional labels and 821,380 non-annotated residues provided an enriched, though potentially noisy, set of nonfunctional positions. Among four supervised machine learning models evaluated, the leaf-wise LightGBM classifier performed best overall, achieving higher average precision and most effectively prioritized annotated active site residues (**fig. S1**) (*11*). Model retraining over 100 randomized resampling iterations maintained high performance, indicating model stability and robust capture of distinguishing features (**fig. S2**).

The trained model computes a residue-level functionality score, referred to as a TAMALE score. A higher score reflects greater model confidence that a given residue is functionally important. The continuous scoring framework allows for relative ranking of all residues within a protein. For visualization, residues are colored where low ranked residues are grey, moderately ranked residues are white, and high-ranking residues are purple (**Fig. 1A**). Benchmarking demonstrates that spatial clustering of highly ranked residues is a strong signature for catalytic and other functional sites. TAMALE analyses of 19,528 human proteins are accessible through a convenient web tool at https://tamale.ccr.buffalo.edu/.

### TAMALE transforms variant effect predictions for active site selectivity

TAMALE exhibits strong selectivity for active site residues compared to Missense. As an initial benchmark, we evaluated the GAPDH2 oxidoreductase for active site detection. Conventional pathogenicity scored 52% of GAPDH2 residues as pathogenic and required the top 37 ranked positions to capture all annotated active site residues. On the other hand, TAMALE identified all three annotated active site residues within the top four ranked positions while prioritizing only 4% of total residues (**Fig. 1B-C**) (*12*). Other highly ranked residues were enriched for ligand binding, indicating the capture of functional signals extending beyond catalysis (**Fig. 1D**).

We evaluated TAMALE on 453 characterized enzymes excluded from model training. The model effectively separated annotated active site residues from the background (**Fig. 1E**). Annotated active site residues were often enriched among positive scores (0 to 5) with meaningful discrimination down to -5. Non-annotated residues were strongly downweighted and predominantly ranged from -5 to -10, with zero representing the model baseline. TAMALE achieved a receiver operating characteristic area under the curve of 0.991 compared to 0.912 for conventional Missense (*P*= < 0.0001, 10,000 bootstrap iterations), demonstrating improved discrimination (**Fig. 1F**). Despite extreme class imbalance (0.26% annotated active site residues), TAMALE achieved substantially higher precision than pathogenicity score alone (**Fig. 1G**). Together, this analysis demonstrates that TAMALE transforms diffuse variant effect signals into highly selective residue-level functional predictions, offering an effective framework for guiding enzymology.

### Structural context attenuates nonspecific signals in variant effect predictions

TAMALE predictions are driven primarily by variant effects and structural features (**Fig. 2A**). To contextualize the sensitivity of a given amino acid position, TAMALE computes an effect score to normalize the residue-level mean pathogenicity to the overall sensitivity of the protein (**Fig. 2B**). Additional effect-based metrics incorporate neighboring primary sequence and spatially proximal residues to capture putative short linear motifs and clustered microenvironments. To distinguish sensitive positions critical for structural integrity, TAMALE incorporates structural features derived from AlphaFold predictions and PAE estimates. Structural residues are often enriched in densely packed cores, whereas functional residues tend to occupy spatially organized and dynamically permissive regions (*13, 14*). Consistently, weighted contact number, which is a measure of local residue packing density and dynamics, is a strong contributor to model performance (**Fig. 2C, fig. S3**). The model also integrates relative solvent accessibility and PAE estimates to capture geometric and conformational constraints (**Fig. 2D-E**). Physicochemical properties modestly contributed to the model, helping to identify structural positions based on hydrophobicity (**fig. S3**). Ablation of individual features resulted in a minor reduction in performance, indicating that TAMALE predictions arise from the combined contribution of multiple Missense-derived and structure-informed signals (**fig. S4**).

**Fig. 2.**
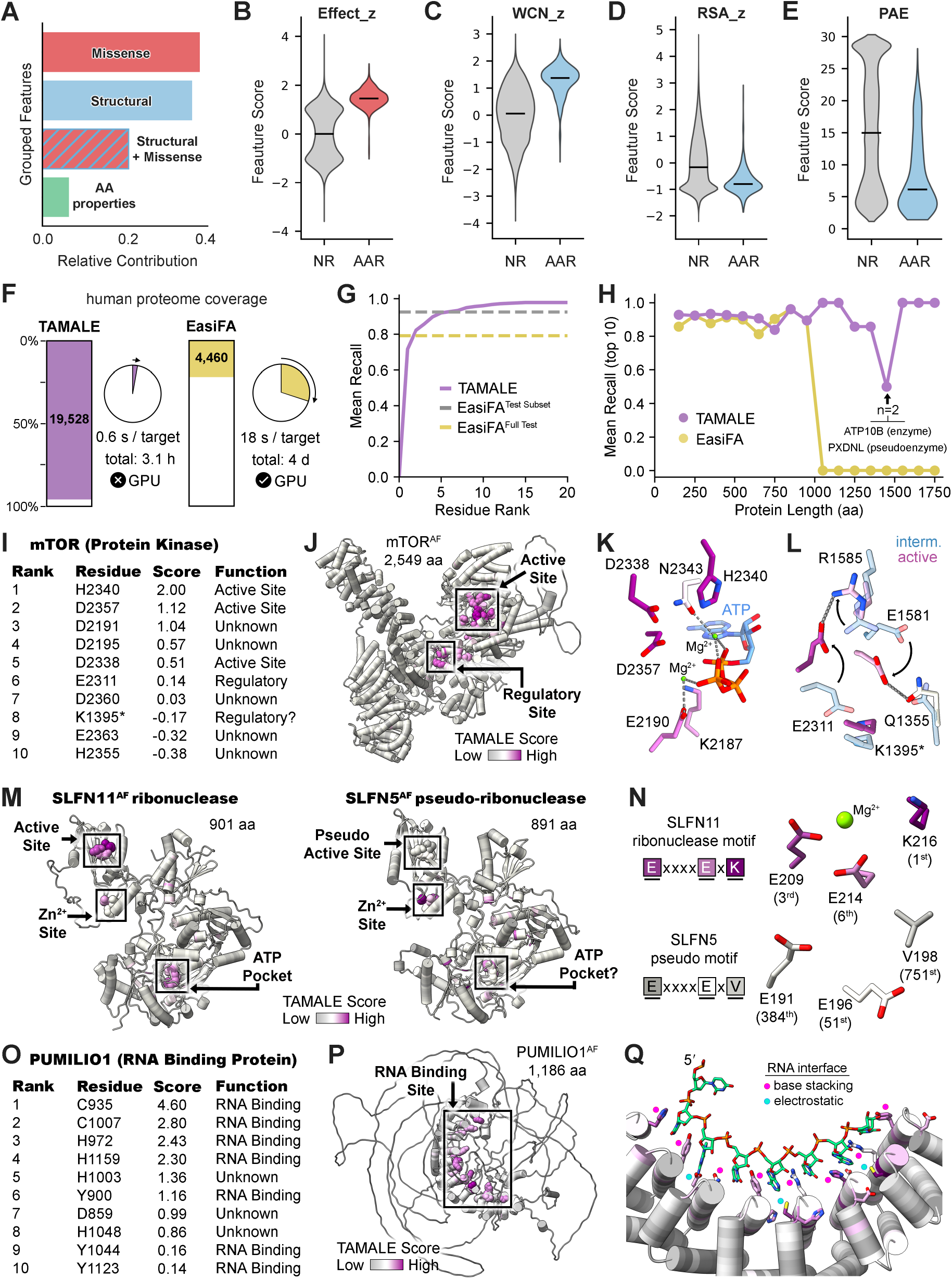
Accurate detection of catalytic and non-enzymatic features by TAMALE. (**A**) Grouped features of TAMALE. (**B-E**) Violin plots of select features between annotated active site residues (AAR) and non-annotated residues (NR). (**F**) Proteome coverage of TAMALE (purple) and high-accuracy mode of EasiFA (yellow, (*6*)) alongside their average target run time and proteome-wide run time (total). (**G**) Mean recall of TAMALE (purple) and EasiFA, reported for the full test set (yellow) and the subset of jobs that completed (grey). (**H**) Recall across protein length. TAMALE correctly discriminated the PXDNL pseudoenzyme from the test set (*58, 59*). (**I**) TAMALE-predicted mTOR functional residues. (**J**) mTOR AF model prediction (*1*). Boxes mark the catalytic and regulatory sites. (**K**) ATP-bound mTOR catalytic site (PDB ID 9ED7; (*23*)). (**L**) mTOR regulatory site overlay comprised of the active state (TAMALE color) and intermediate state (interm., blue) (PDB ID 9ED7, 9ED8, (*23*)). Asterisk marks an unreported functional residue. (**M**) SLFN11 and SLFN5 AF model predictions (*1*). (**N**) SLFN11 ribonuclease motif and SLFN5 pseudomotif (PDB ID 7PPJ; 9ERD (*29, 30*)). (**O-P**) TAMALE analysis of PUMILIO1 with the RNA binding site boxed. (**Q**) Crystal structure of RNA bound PUMILIO1 (PDB ID 1M8X, (*31*)).

### Proteome-wide functional predictions without EC annotation

We benchmarked TAMALE against EasiFA, a state-of-the-art enzyme annotation tool that integrates a protein language model with enzyme chemistry. In its high-accuracy mode, EasiFA analyzes 22% of the human proteome over four days using GPU infrastructure but requires EC annotation, limiting its analysis to known enzymes of moderate size (< 1,000 residues) (**Fig. 2F-H**) (*6*). In contrast, TAMALE operates on a standard laptop independently of EC annotation and is insensitive to protein length, matching the performance of EC-guided EasiFA while processing 96% of the human proteome in ∼3 hours to generate predictions for 19,528 human proteins spanning both enzymes and non-catalytic proteins (**Fig. 2F, 2H**). Failed runs were primarily attributed to a missing input file, which can often be resolved using AlphaFold3 (*15*).

### Generalizability across enzyme classes

We evaluated the performance of TAMALE across the seven enzyme classes. Without requiring EC labels or reaction information, TAMALE had the highest sensitivity and selectivity for transferases, hydrolases, and translocases and strong performance for oxidoreductases and isomerases (**fig. S5**). A minor reduction in performance for lyases can be partly attributable to their underrepresentation in model training (1.8%) and a tendency to prioritize binding over catalytic residues (**fig. S6**). A modest decrease in recall was also observed for ligases (**fig. S5**), consistent with the reduced prioritization of lysine residues, which frequently serve catalytic roles in this enzyme class (**fig. S7**) (*16*). For example, in DNA ligase I, the catalytic lysine residue is ranked below two active site residues (**fig. S8**) (*17*).

Across enzyme classes, annotated active site residues are typically ranked first, with additional unannotated active site residues within the top ten ranked positions. Exceptions to this trend often reflected true functional signal. In TOP2B, the annotated active site residue ranks second while the top ranked residue (D562) is an unannotated residue coordinating a catalytic magnesium ion (*18*). In DUSP28, the annotated catalytic residue (C103) ranks fourth while the top three residues are unannotated ligand binding residues (**fig. S8**) (*19*). Similarly, in MRE11, the annotated active site residue is second, while the remaining top five ranked residues contribute to the catalytic binding environment (**fig. S8**) (*20*). Systematic analysis of annotated binding residues demonstrates that TAMALE prioritizes functionally important positions beyond active site residues, suggesting the current benchmarks may underestimate its full predictive capacity (**fig. S6**).

### TAMALE effectively detects functional sites within large proteins

To determine whether TAMALE can identify diverse functional features in large, structurally complex enzymes, we analyzed the 2,549-residue kinase mTOR responsible for cellular homeostasis and metabolism. Highly ranked residues cluster to form two predicted functional sites (**Fig. 2I-J**). The top-scoring site is located within the kinase domain, where TAMALE identifies residues forming the catalytic center. Key residues include H2340 (1^st^) which contacts both the substrate and ATP to stabilize the transition state, D2357 (2^nd^) and N2343 (56^th^), which coordinate catalytic magnesium ions, and D2338 (5^th^), which positions the protein substrate for nucleophilic attack (**Fig. 2K**) (*21*). K2187 and E2190 (top 15) also form a salt bridge that supports ATP binding (**Fig. 2K**) (*22*). The second cluster aligns to a known regulatory site between the kinase and FAT domains, capturing electrostatic interactions associated with its active state, including E2311 (6^th^) and R1585 (35^th^) (**Fig. 2L**) (*23*). Interestingly, FAT residue K1395 (8^th^) remains uncharacterized, but lies within the regulatory site and may exert an unreported role in mTOR control. TAMALE also retained high performance on an extreme case, accurately identifying catalytic and substrate binding residues in the 3,546-residue USP34 deubiquitinating enzyme (**fig. S9**, see supplementary text).

### Evaluating functional features of enzyme scaffolds

The BRCA1-BARD1 complex is a scaffold for nucleosome ubiquitination by E2 enzymes, such as UBE2D3. TAMALE correctly ranks the catalytic cysteine of UBE2D3 (C85) as the top ranked position, with active site residues D87 and D117 in the second and fourth positions, respectively (**fig. S10**) (*24*). It further enriches E1 (e.g. R5, E9) and E3 interacting residues, indicating recognition of protein-protein interaction interfaces (*25*). Within BRCA1-BARD1, TAMALE prioritized scaffolding features, such as zinc finger motifs and phosphopeptide binding sites (**fig. S10**) (*24, 26*). Performance is reduced in low-complexity regions, consistent with Missense derived features and defines a limitation of TAMALE applicability (*9*).

### TAMALE differentiates pseudoenzymes from active enzymes

To test whether TAMALE distinguishes pseudoenzymes from catalytic homologs, we analyzed the Schlafen (SLFN) ribonuclease family (*27*). The catalytic member SLFN11 harbors a consensus <u>E</u>xxxx<u>E</u>x<u>K</u> ribonuclease motif, whereas the inactive member SLFN5 carries a degenerate <u>E</u>xxxx<u>E</u>xV motif (*28, 29*). Despite high structural homology and sequence conservation, TAMALE correctly prioritizes SLFN11 catalytic residues, ranking E209 3^rd^, E214 6^th^ and K216 1^st^, while deprioritizing SLFN5’s corresponding pseudomotif (E191 384^th^, E196 51^st^, V198 751^st^) (**Fig. 2M-N**) (*28*). P-loop and Walker B motifs are also prioritized, suggesting potential function to the cryptic ATPase site in both SLFN11 and SLFN5 (**fig. S11**) (*29, 30*). Collectively, these results show TAMALE can discriminate functional elements between closely related protein family members.

### Recognition of nucleic acid binding residues

To assess whether TAMALE identifies nucleic acid binding residues, we analyzed PUMILIO1, an RNA binding protein involved in post-transcriptional gene regulation. TAMALE strongly enriched for residues known to form the concave RNA interaction surface, including canonical motif positions involved in RNA recognition (**Fig. 2O-Q**) (*31*). Notably, it identifies residues at motif position 13 that mediate nucleobase stacking and polar residues at motif positions 12 and 16 that contribute to electrostatic RNA interactions. This analysis highlights the potential of leveraging TAMALE to characterize other nucleic acid binding sites.

### Identification of allosteric sites in signal-transduction proteins

We then assessed the model’s performance on KRAS, a GTPase molecular switch, and a high-value therapeutic target (*32*). TAMALE identified three discrete clusters corresponding to the nucleotide-binding pocket and druggable allosteric sites (**fig. S12**). Within the catalytic site, it correctly prioritized key residues involved in GTP binding and hydrolysis, including P-loop residues (G15, K16, S17) and nucleobase contact residues (N116, K117, D119), as well as Mg^2+^ ion coordination residues and the catalytic residue from the RASA1 GTPase activating protein (**fig. S12**) (*33, 34*). It also identifies known allosteric pockets (SI/II and S-II) and switch regions involved in effector binding and signaling, including contacts with the RAF proto-oncogene (**fig. S12**) (*33, 35, 36*). We also tested the non-enzymatic ADRB2 GPCR for the accurate identification of multiple allosteric sites (**fig. S13**, see supplementary text) (*37-41*). These analyses reveal the capacity for identifying allosteric sites in signaling proteins.

### *De novo* identification of enzymatic signatures

To evaluate whether TAMALE can nominate new enzyme function, we examined whether score distributions distinguish enzymes from non-catalytic proteins. Enzymes exhibit a broader score range (10-20) compared to non-enzymatic proteins (5-15), with partial overlap that may include proteins with unrecognized catalytic activity (**Fig. 3A**). For example, mTOR kinase has a similar score distribution to the uncharacterized protein C5orf22 (**Fig. 3A**). High scoring residues in C5orf22 cluster into a putative active site resembling the validated Arg1 catalytic center of the ureohydrolase superfamily (**Fig. 3B-C**) (*42*). In Arg1, higher TAMALE scores align with canonical positions of the arginase motifs, providing a residue-level readout of functional relevance (**Fig. 3D**). C5orf22 shows a similar signature, including a high scoring extended DxHxD motif, supporting its proposed role as an atypical peptide-specific arginase (**Fig. 3D**) (*43*).

**Fig. 3.**
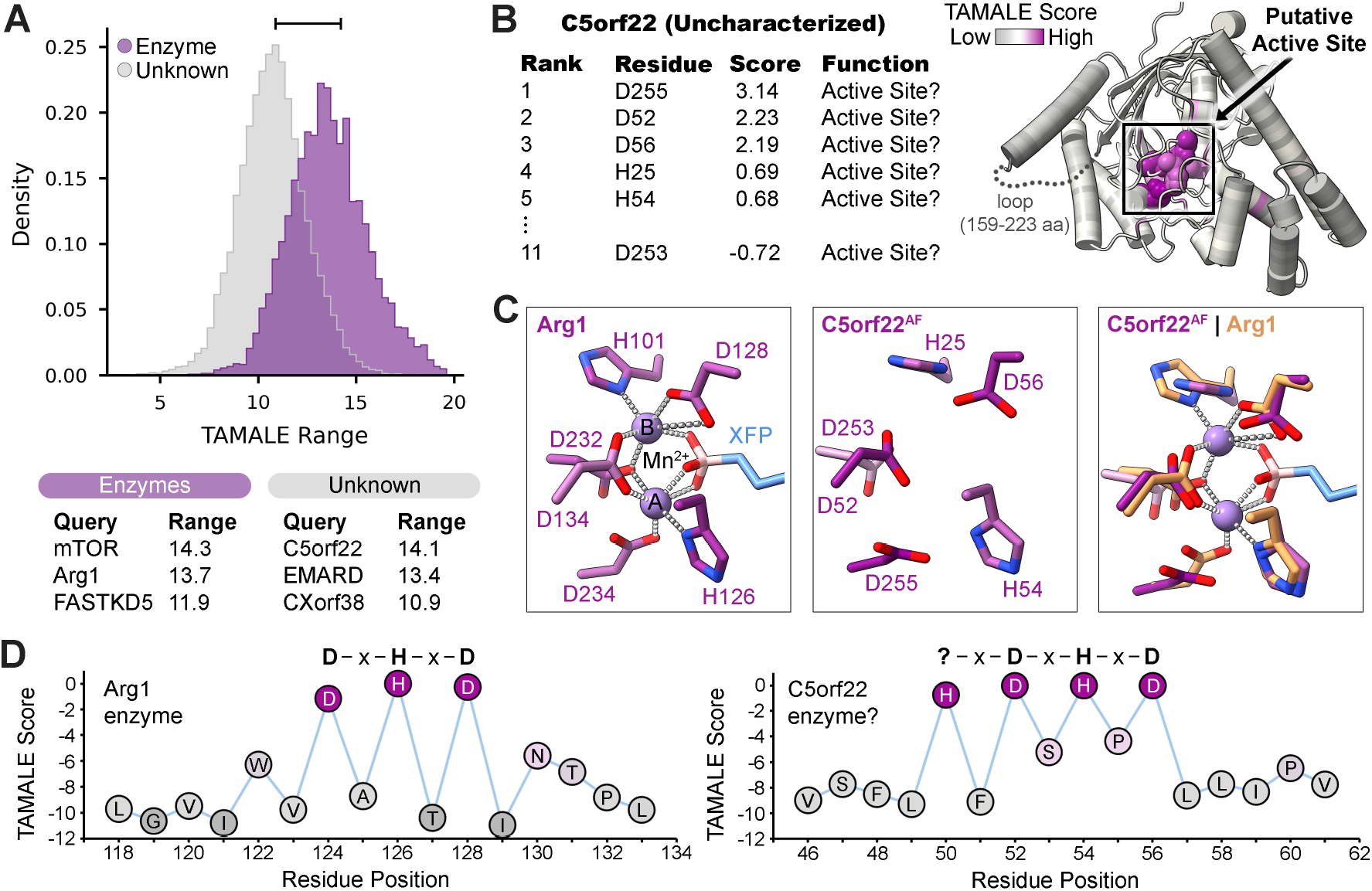
TAMALE enables *de novo* proteome-wide analysis. (**A**) Distribution of the TAMALE score range relative to the normalized density of annotated enzymes (purple, Enzyme) and proteins with no known annotated active site (grey, Unknown). The enzyme group comprises 4,470 proteins and the unknown group is a total of 15,059 proteins. Horizontal bracket marks the overlapping region where proteins with unrecognized catalytic activity may be enriched. (**B**) Top ranked C5orf22 residues along with the AlphaFold (AF) model prediction colored by TAMALE score (*1*). Box marks a cluster of high ranking residues forming a putative enzyme active site. (**C**) Human Arg1 active site alongside the putative C5orf22 active site with TAMALE score colors (PDB ID 7KLK; (*60*)). A model overlay of Arg1 (orange) and C5orf22 demonstrate the predicted structural resemblance. Manganese ions (Mn^2+^) are pale violet, the Arg1 inhibitor compound 3a (XFP) is shown in blue, and dotted lines define metal ion coordination. (**D**) TAMALE scores for the consensus arginase motif DxHxD from validated arginase Arg1 and putative enzyme C5orf22.

### Hypotheses generation for emerging HEPN ribonucleases

HEPN ribonucleases are conserved across evolution and function as effector components in systems such as toxin-antitoxin and CRISPR-Cas modules (*44*). They are defined by a consensus RϕxxxH catalytic motif (where ϕ is D, N, or H and x is any amino acid) and, in some cases, an additional ExxxR/K motif (*45*). Catalysis requires dimerization, which forms a composite active site from juxtaposed RϕxxxH motifs (*44*). TAMALE accurately identified all three active site residues (R1, ϕ2, and H6 of <u>Rϕ</u>xxx<u>H</u>) in Las1L (**fig. S14**) (*44, 46*), supporting its utility for characterizing hypothetical HEPN proteins. For EMARD, TAMALE prioritizes the putative ^134^<u>E</u>RALG and ^186^<u>RN</u>VLW<u>H</u> motifs consistent with prior predictions (**Fig. 4**) (*45, 47*). Importantly, it also revealed a second unreported putative catalytic domain containing ^299^<u>E</u>TGLR and ^369^<u>RD</u>HLS<u>H</u> motifs, suggesting EMARD may adopt an asymmetric intramolecular composite active site analogous to CRISPR-Cas13 (**Fig. 4C**) (*48*). If validated, this would represent the first intramolecular HEPN enzyme in humans. For CXorf38, TAMALE identifies the previously proposed ^162^<u>RN</u>EIM<u>H</u> catalytic motif, but reassigns catalytic importance from the proposed acidic residue of ^39^<u>E</u>VLSF to D134 (1^st^) (**Fig. 4E-G**) (*45*). Structural modeling suggests the active site is partially occluded by a downstream helical bundle, consistent with a possible autoinhibited state analogous to the bacterial toxin-antitoxin systems (**Fig. 4G**) (*49*). A truncated dimer model restores a canonical HEPN arrangement centered on the proposed H167–D134 dyad (**Fig. 4H-I**). Taken together, the algorithm can propose mechanistic hypotheses in sequence divergent superfamilies.

**Fig. 4.**
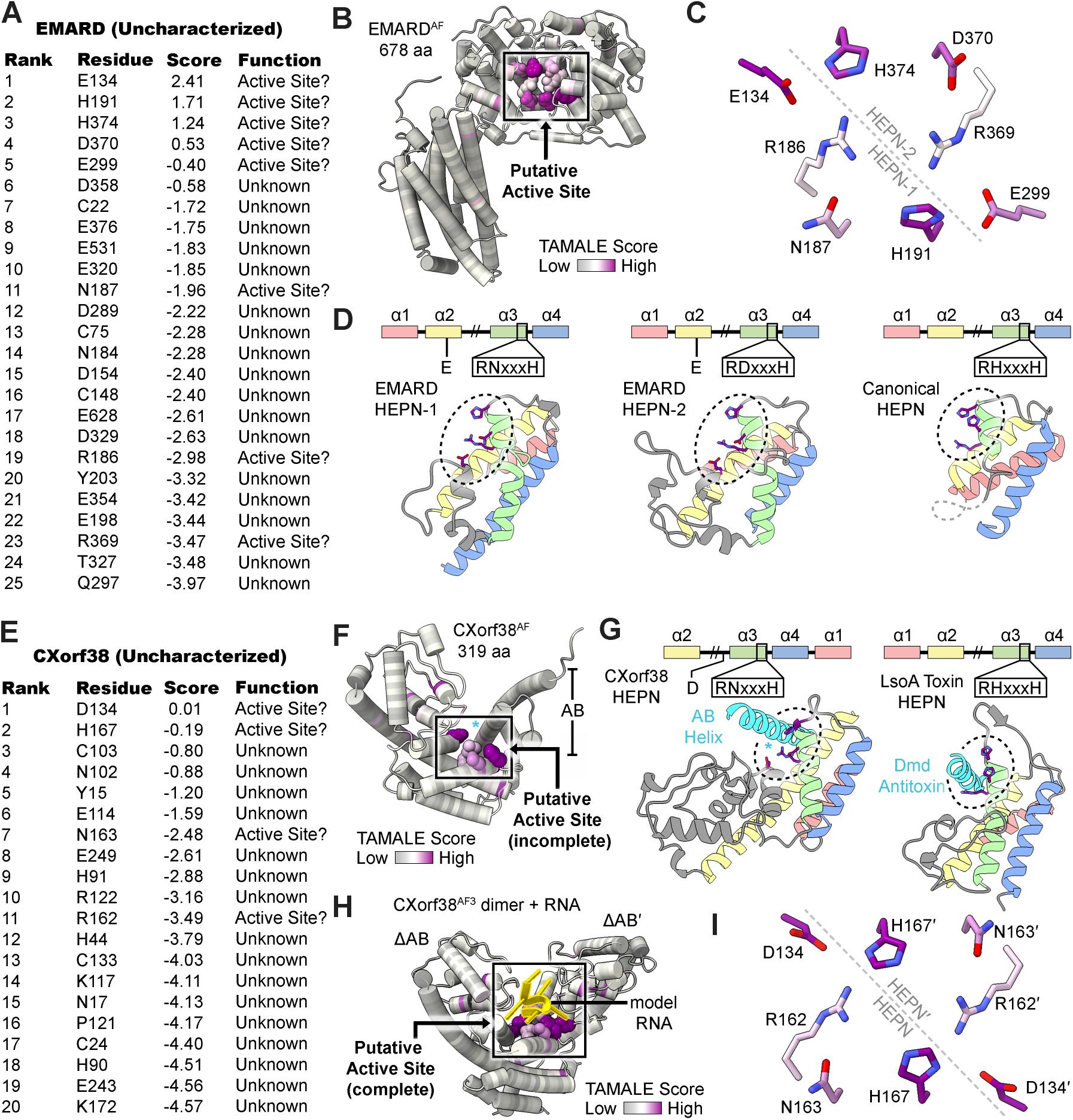
Molecular insights revealed by predicting HEPN active site residues. (**A**) TAMALE-predicted EMARD functional residues. (**B**) EMARD AlphaFold (AF) model prediction with TAMALE coloring revealing a cluster representing a putative intramolecular HEPN-1–HEPN-2 active site (boxed) (*1*). (**C**) Predicted asymmetric architecture of the composite EMARD HEPN site. (**D**) Predicted arrangement of high ranking EMARD residues is reminiscent to the canonical and validated Las1L HEPN, lending support that EMARD is a HEPN enzyme. (**E**) TAMALE-predicted CXorf38 functional residues. (**F**) CXorf38 AF model prediction with TAMALE coloring reveals a cluster representing a putative HEPN active site (boxed) (*1*). (**G**) Predicted arrangement of high ranking CXorf38 residues is similar to the HEPN-encoded bacterial toxin-antitoxin system (PDB ID 5HY3; (*49*)), suggesting CXorf38 may adopt an autoinhibited (incomplete) HEPN state. (**H**) Prediction of the CXorf38 antitoxin-like C-terminal truncation reveals a homodimer that threads RNA along the putative active site (*15*). (**I**) Inset of the predicted CXorf38 dimer proposes a canonical (complete) composite HEPN arrangement with the proposed His-Asp dyad. Cyan asterisk marks the antitoxin-like helix.

### TAMALE discovers an atypical catalytic site in FASTKD5

We applied TAMALE to the understudied mitochondrial ribonuclease FASTKD5, which processes specific mitochondrial RNA junctions including ATP8/6-CO3 (**Fig. 5A**) (*50*). Although its catalytic activity has been reconstituted *in vitro* and mutagenesis of conserved residues has identified sensitive amino acid positions, the composition of its catalytic center remains unresolved (*50*). Structural homology to the metal-dependent PD-DxK nuclease superfamily has been proposed but is inconsistent with FASTKD5’s metal-independent ribonuclease activity (**Fig. 5B-C**) (*51*). Since existing computational tools fail to define a FASTKD5 active site, we applied TAMALE to nominate candidate catalytic residues (**Fig. 5D**). TAMALE predicts a compact functional site within the poorly characterized FASTKD5 helix-turn-helix region. The top ranked residue, H395, lies within a positively charged concave surface and clusters with K428 (2^nd^) and D429 (10^th^), all previously shown to be vital for mitochondrial RNA processing (**Fig. 5B**) (*50*). Additional nearby residues E470 (17^th^) and H471 (9^th^) extend this site. In contrast, the PD-DxK-like residue D589 (7^th^) is not strongly prioritized and does not cluster with other high-ranking residues, leaving its functional role unclear (**Fig. 5D**). Collectively, TAMALE generates a testable hypothesis for FASTKD5 ribonuclease activity.

**Fig. 5.**
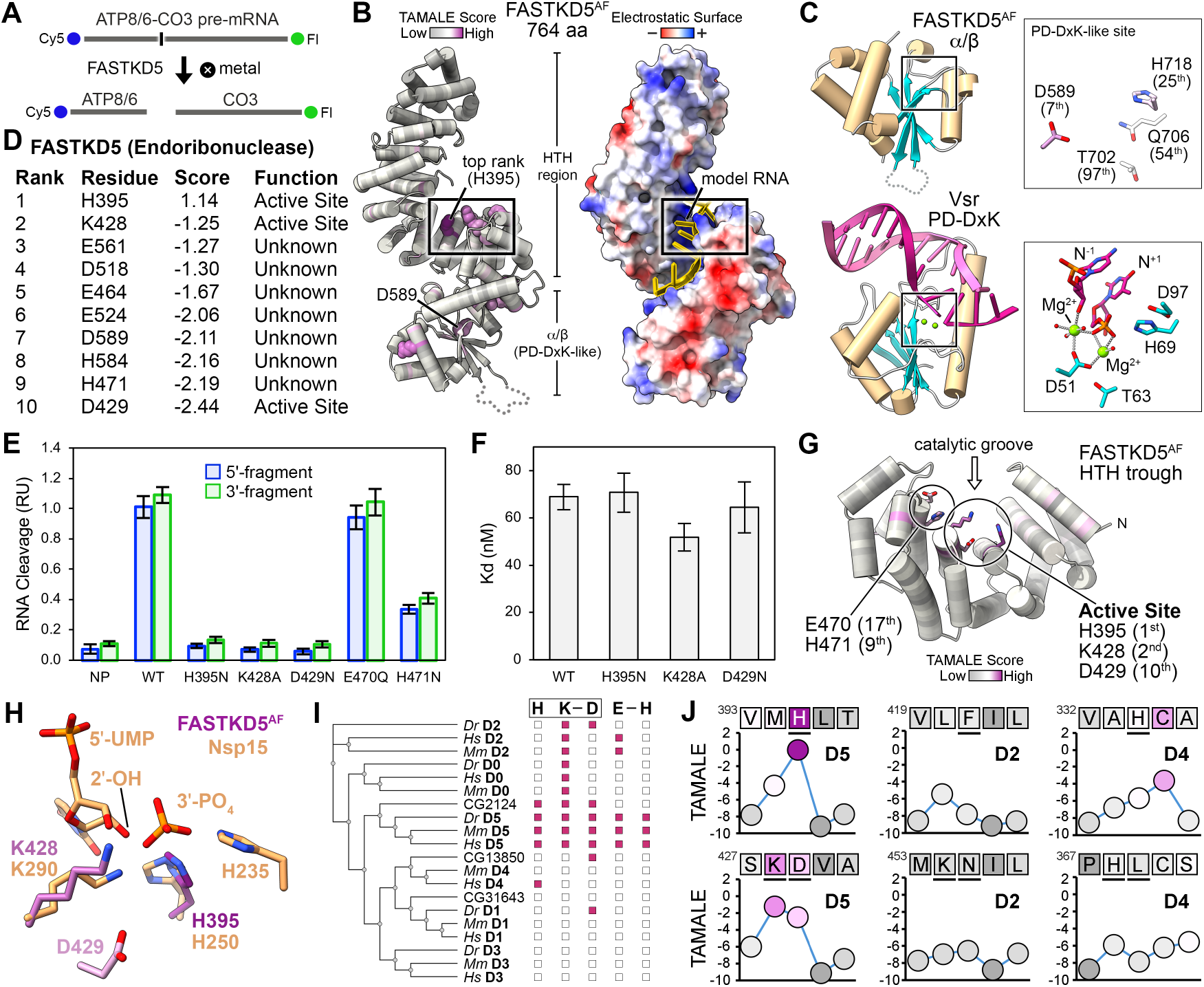
Discovery of the unconventional FASTKD5 catalytic center. (**A**) Schematic of mitochondrial precursor mRNA (pre-mRNA) cleavage by the metal-independent FASTKD5 ribonuclease. Blue and green dots represent the 5’-Cy5 and 3’-fluorescein (Fl) labels used in this study. (**B**) FASTKD5 AlphaFold (AF) model prediction (*15*). Surface representation displays electrostatic surface potentials, showing the putative RNA-binding surface. The extended loop (residues 594-694) is displayed as a dotted line. (**C**) FASTKD5 α/β fold next to the crystal structure of the *Escherichia coli* DNA-bound Vsr PD-DxK endonuclease (PDB ID 1CW0; (*61*)). Black box marks the validated Vsr endonuclease site and the equivalent FASTKD5 site. (**D**) Top TAMALE ranked FASTKD5 residues. (**E**) *In vitro* RNA cleavage activity of FASTKD5 variants and a no protein control (NP). The bar defines the mean and error bars represent the standard deviation from three independent replicates. (**F**) *In vitro* RNA binding activity of select FASTKD5 variants where the bar represents the mean dissociation constant (Kd) and error bars are the standard deviation from two independent replicates. (**G**) Proposed architecture of the FASTKD5 catalytic trough. (**H**) Overlay of the FASTKD5 active site residues with ribonuclease Nsp15 (PDB ID 7K0R; (*54*)). (**I**) Phylogenetic tree of FASTK family members from *Homo sapiens* (Hs), *Mus musculus* (Mm), *Danio rerio* (Dr), and *Drosophila melanogaster* (CG numbers; (*62*)) (*2, 63, 64*). Mitochondrial isoform of FASTK is D0 and FASTKD1-D5 are D1-D5, respectively. Evolutionary conservation of equivalent residues to H395, K428, and D429 (H K-D) and proximal E470 and H471 residues (E-H) are marked by a pink box. (**J**) TAMALE scores of FASTKD5 and select non-catalytic family members (D2 and D4) with equivalent residues underlined.

We purified monomeric human FASTKD5 protein and confirmed its suitability for biochemical analysis (**fig. S15**). Recombinant FASTKD5 cleaves a model ATP8/6-CO3 RNA junction *in vitro*, generating the expected 5′- and 3′-fragments (**Fig. 5A, fig. S15**) (*50, 52*). To characterize the function of the TAMALE site, we executed a mutagenesis campaign where histidine residues were mutated to asparagine (H395N, H471N) to prevent general acid-base chemistry, aspartate was replaced with asparagine (D429N) and glutamate was changed to glutamine (E470Q) to remove the negative charge, and lysine was mutated to alanine (K428A) to remove the positive charge and prevent hydrogen bonding (**fig. S15**). E470Q retained wild-type RNA cleavage and binding activity, whereas H471N reduced RNA cleavage due to slower RNA association (**Fig. 5E, fig. S15**). Conversely, H395N, K428A, and D429N substitutions abolish RNA cleavage activity, consistent with previous reports. (**Fig. 5E**) (*50*). H395N preserved RNA binding affinity and kinetics supporting a direct catalytic role, while K428A and D429N altered early RNA binding events (**Fig. 5F, fig. S15**). These data define a functional His-Lys-Asp ribonuclease motif and confirm accurate discovery of a FASTKD5 catalytic residue.

The FASTKD5 His-Lys-Asp ribonuclease motif forms a compact active site at the base of a predicted ∼15 × 15 × 17 Å trough resembling non-enzymatic scaffolds from the Trypanosomal RNA editing complexes (**Fig. 5G**) (*1, 53*). To our knowledge, this represents the first assignment of ribonuclease activity to this predicted helix-turn-helix fold. Although it is structurally distinct from SARS-CoV-2 Nsp15, FASTKD5 shares parallels where H395 and K428 overlap with Nsp15 H250 and K290, consistent with a role in promoting nucleophilic attack and transition-state stabilization, respectively (**Fig. 5H**) (*54*). And yet, FASTKD5 lacks the second catalytic histidine found in Nsp15 but instead positions D429 adjacent to H395, which may modulate catalysis through a water-mediated interaction (*55, 56*). Consistent with a functional signature, the His-Lys-Asp motif is conserved within the FASTKD5 clade but absent in structurally similar non-catalytic family members and is present in *Drosophila melanogaster* CG2124, suggesting an unrecognized FASTK ribonuclease (**Fig. 5I-J**).

## Conclusions

TAMALE provides a scalable framework for identifying residues that directly contribute to catalysis, molecular interactions, and regulation without requiring prior functional annotation. By repurposing missense variant effect predictions as a form of *in silico* saturating mutagenesis, TAMALE demonstrates that population-derived sequence constraints encode rich and underexploited information on protein function. This strategy reimagines variant effect predictions not simply as a measure of pathogenicity, but as an intermediate resource of latent functional signals that can be mined for mechanistic insight across the proteome.

By leveraging diffuse biochemical constraints embedded in population variation, TAMALE minimizes the reliance on human curated knowledge thereby reducing bias towards characterized functional sites. Rather than primarily enriching for known functional signatures, this model emphasizes downweighting of structural signals to allow functional patterns to emerge in a manner that is largely independent of prior functional knowledge. TAMALE’s capacity to identify new function was demonstrated with FASTKD5, where TAMALE ranked the catalytic histidine residue in the top position despite the lack of structural homology to known nucleases. Although applied here to the human proteome, this framework is organism-agnostic and can be applied broadly as variant effect resources expand across evolutionary space.

## Acknowledgments

We are grateful to Dr. Olivia A. Fraser for assistance in maintaining insect cells. We thank Dr. Thomas Grant and Dr. Andrew Gulick for their critical reading of this manuscript. We are grateful to Dr. Thomas Westbrook for providing lab space and Dr. Alex Vecchio for access to the biolayer interferometry instrumentation. Support provided by the Center for Computational Research at the University at Buffalo (*57*).

## Funding

This work was supported by the US National Institutes of Health grant R35GM147123 (MCP) issued to Baylor College of Medicine and transferred to the State University of New York at Buffalo.

## Author contributions

Conceptualization: JVR, MCP

Methodology: JVR, BJC, MCP

Investigation: JVR, BJC, MCP

Visualization: JVR, BJC, MCP

Funding acquisition: MCP

Project administration: MCP

Supervision: MCP

Writing – original draft: JVR, MCP

Writing – review & editing: JVR, BJC, MCP

## Competing interests

MCP and JVR are inventors on a pending patent application related to the technology described in this work. The remaining author declares no competing interests.

## Data, code, and materials availability

For broad and rapid adoption of TAMALE data, output files for 19,528 human proteins are available using the TAMALE web tool found at https://tamale.ccr.buffalo.edu/. The TAMALE code was deposited to GitHub at https://github.com/PillonLab/TAMALE. © 2026 Research Foundation for the State University of New York & Baylor College of Medicine. All rights reserved. The TAMALE software and all output data from TAMALE software, including derivatives and Adapted Material, are made available for non-commercial research use by non-profit entities only under the Creative Commons Attribution-Non-Commercial-ShareAlike 4.0 (CC BY-NC-SA 4.0) license. Patent applied for. To discuss commercial licensing, please contact. Inputs used by TAMALE were obtained through the AlphaFold database found at https://alphafold.ebi.ac.uk/. EasiFA analysis was found at https://cadd.zju.edu.cn/easifa/.

## Supplementary Materials

Materials and Methods

Supplementary Text

Figs. S1 to S15

Tables S1 to S3

References (*65*–*85*)

Data S1

## Notes

https://tamale.ccr.buffalo.edu/

