## Supplementary Materials for "Accurate Identification of Functional Residues Across the Human Proteome with TAMALE"

**The PDF file includes:**

Materials and Methods
Supplementary Text
Figs. S1 to S15
Tables S1 to S3
References

**Other Supplementary Materials for this manuscript include the following:**

Data S1

### Materials and Methods

Data Curation for Pipeline

Using a target's UniProt accession number, three publicly available output files previously derived and accessible from the AlphaFold Database (https://alphafold.ebi.ac.uk, (*1, 9*)) are imported for the pipeline: the AlphaFold Missense data (.csv), the consensus AlphaFold predicted model (.mmCIF), and the associated predicted aligned error (PAE) (.json).

Gathering and Computing Residue Attributes and Statistics

A comprehensive feature table is generated using the imported data, resulting in 48 unique residue-specific metrics that can be generally categorized as i) structure features, ii) Missense-derived features, and iii) physicochemical features.

*Initial structure features:* We parse the mmCIF file to build a residue index structure table using chain A of the AlphaFold model, storing the residue's position number (1-n), one-letter amino acid code, and C-alpha atom coordinates (x, y, z). We compute the weighted contact number for each residue ($\mathrm{WCN}_i$) using the equation:

$$\mathrm{WCN}_i = \sum_{\mathrm{j} \neq \mathrm{i}} \frac{1}{\| r_i - r_j \|^2}$$

where $i$= indexed position, $j$= all other positions, and $r$= C-alpha coordinate vector (x, y, z) (*65*). We calculate each residue's solvent accessibility surface area (ASA) using the Shrake-Rupley algorithm (*66*) and then compute a relative solvent accessibility (RSA) to normalize against that amino acid using the following equation:

$$RSA\_rel_i = \frac{ASA_i}{MAX\_ASA_{(a_i)}}$$

where $RSA\_rel_i$ = normalized RSA for position $i$, $ASA_i$= Shrake-Rupley accessible surface area for position $i$, and $MAX\_ASA_{(ai)}$= maximum accessible surface area for that amino acid (a) at position $i$. We define the maximum accessible surface area (Å$^2$) for amino acid (a) as 106 (Ala), 248 (Arg), 157 (Asn), 163 (Asp), 135 (Cys), 198 (Gln), 194 (Glu), 84 (Gly), 184 (His), 169 (Ile), 164 (Leu), 205 (Lys), 188 (Met), 197 (Phe), 136 (Pro), 130 (Ser), 142 (Thr), 227 (Trp), 222 (Tyr), and 142 (Val) (*67*). Using the PAE json file, we parse the N x N matrix ($\mathrm{PAE}_{\mathrm{i,j}}$) to compute a single-residue PAE by taking the mean, defined as the average aligned error of that position compared to all other positions as shown below:

$$pae_i = \frac{1}{N} \sum_{j=1}^{N} PAE_{i,j}$$

where $pae_i$= mean predicted aligned error of residue $i$, $i$= index of residue in the protein, j= index of any other residue of the protein, $pae_{i,j}$= predicted aligned error of residue $i$ when the structure is aligned on residue j, and $N$= total number of residues in the protein (*1*).

*Missense-derived features:* The “protein_variant” and “am_pathogenicity” columns were parsed from the Missense csv file, building a table comprising the residue position ($i$), wild type one-letter amino acid code at position $i$ ($a_i$), mutant one-letter amino acid code (b), and the missense pathogenicity score value (s or cost) (*9*). Next, we compute the mean cost of each residue position ($i$) alongside the mean cost of all remaining positions within the protein, excluding position $i$, to determine the background cost. We define the set of all missense variants for each position ($i$) as $S_i$ where $S_i = \{S_{i,1}, S_{i,2}, S_{i,3} \ldots S_{i,n}\}$. We compute a mean missense score for each position ($\mu_i$) and store this value as ‘pos_mean’ using the following equation:

$$\mu_i = \frac{1}{n_i} \sum_{k=1}^{n_i} s_{i,k}$$

where $n_i$= number of variants at position $i$, $k$= index in the set $S_i$, and $S_{i,k}$= the pathogenicity score of the $k^{\text{th}}$ mutation at position $i$. We also compute a background mean missense score for each position ($\mu_{rest,i}$) and store this metric as ‘rest_mean’ using the equation below:

$$\mu_{rest,i} = \frac{1}{n_{rest,i}} \sum_{k=1}^{n_{rest,i}} s_{rest,i,k}$$

where $n_{rest,i}$= number of variants in the entire protein excluding position $i$, $k$= index in the set $S_{rest,i}$, and $S_{rest,i,k}$= the cost of the $k^{\text{th}}$ mutation in the protein excluding position $i$. Furthermore, we compute the variance of cost for each residue position ($i$) alongside the variance of cost for all remaining positions within the protein, excluding position $i$. Variance was calculated using Bessel’s correction (division by $n_i - 1$) to obtain an unbiased estimate of variance (*68*). Variance in pathogenicity (or cost) for each position ($Var_i$) was calculated using the equation:

$$Var_i = \frac{1}{n_i - 1} \sum_{k=1}^{n_i} (s_{i,k} - \mu_i)^2$$

whereas variance of the background pathogenicity ($Var_{rest,i}$) was determined using:

$$Var_{rest,i} = \frac{1}{n_{rest,i} - 1} \sum_{k=1}^{n_{rest,i}} (s_{rest,i,k} - \mu_{rest,i})^2$$

where $n_{rest,i}$= number of variants in the entire protein excluding position $i$, $k$= index in the set $S_{rest,i}$, $S_{rest,i,k}$= the cost of the $k^{\text{th}}$ mutation in the protein excluding position $i$, and $\mu_{rest,i}$= mean cost for all variants in the protein excluding $S_i$. To account for the background pathogenicity at each position, we computed an effect score ($Effect_i$) and stored the value as ‘pos_effect’ using the equation below:

$$Effect_i = \mu_i - \mu_{rest,i}$$

where $\mu_{rest,i}$= mean pathogenicity across the protein excluding position $i$ and $\mu_i$= mean pathogenicity score for position $i$. We performed statistical analyses on each position to compute p and q values for each pathogenicity set. A Welch t-test (*69*) was used to derive metrics such as a t statistic ($t_i$), effective degrees of freedom ($v_i$), and two-sided p value ($p_i$):

t statistic:

$$t_i = \frac{effect_i}{\sqrt{\frac{Var_i}{n_i} + \frac{Var_{rest,i}}{n_{rest,i}}}} = \frac{\mu_i - \mu_{rest,i}}{\sqrt{\frac{Var_i}{n_i} + \frac{Var_{rest,i}}{n_{rest,i}}}}$$

effective degrees of freedom:

$$v_i = (\frac{\frac{Var_i^2}{n_i} + \frac{Var_{rest,i}^2}{n_{rest,i}})^2}{\frac{\left(\frac{Var_i^2}{n_i}\right)^2}{n_i - 1} + \frac{\left(\frac{Var_{rest,i}^2}{n_{rest,i}}\right)^2}{n_{rest,i} - 1}}$$

two-sided p value:

$$p_i = 2\left(1 - F_{t,v_i}(|t_i|)\right)$$

We applied the Benjamini-Hochberg correction to control for the false discovery rate (*70*). First, the compiled set of p-values from all positions were defined as $\{ p_1, p_2, \ldots . p_n \}$. Second, p-values were ranked in ascending order $p_i(1) \leq p_i(2) \leq p_i(3) \leq \cdots \leq p_i(n)$ where $(k)$ is the rank of that p-value (1= smallest value). Third, we computed the Benjamini-Hochberg adjusted value ($q_{(k)}$) for each p-value:

$$q_{(k)} = \frac{m}{k}\, p_{(k)}$$

where $m$ is the total number of statistical tests (or p-values) in the set. Finally, we implement monotonicity ($q_{(k)}^{monotone}$) to retain the original p-value ranking using the equation:

$$q_{(k)}^{monotone} = \min_{t \geq k} q_{(t)}$$

where $t \geq k$= all ranks from $k$ to the largest rank ($m$) and min$q_{(t)}$= the smallest value of the Benjamini-Hochberg adjusted p-values (*71*). We apply the following thresholds to define a statistically significant position from other positions where i) before Benjamini-Hochberg adjustment: $sig_{p05_i} = True\ \ if\ q_{(i)}^{monotone} \leq 0.05; else\ False$, ii) after Benjamini-Hochberg adjustment: $sig_{q05_i} = True\ \ if\ q_{(i)}^{monotone} \leq 0.05; else\ False$, and iii) after Benjamini-Hochberg adjustment and the position has an effect score > 0 (suggesting functional importance): $sig_{q05_{pos_i}} = True\ \ if\ (q_{(i)}^{monotone} \leq 0.05)\ and\ (effect_i > 0\ ); else\ False$. We marked significant residues cleanly for downstream functions by $sig_i = sig_{q05_{pos_i}}$. We also carried forward the extreme minimum (pos_min) and maximum (pos_max) missense pathogenicity score for each position along with their associated amino acid mutation (min/max_cost_aa) as defined by pos_min= minimum pathogenicity score for position $i$ in set $S_i$, min_cost_aa= one-letter amino acid code for $k$ in $S_i$ with minimum value, pos_max= maximum pathogenicity score for position $i$ in set $S_i$, and max_cost_aa= one-letter amino acid code for $k$ in $S_i$ with maximum value. We generated a respective delta for each metric by calculating the difference between the position's extreme value and the mean where min_cost_delta_from_pos_mean$_i$= pos_min$_i$ − $\mu_i$ and max_cost_delta_from_pos_mean$_i$= pos_max$_i$ − $\mu_i$. We described the spread of the pathogenicity scores by calculating the standard deviation, $std_i = \sqrt{Var_i}$. For each position ($i$), we tracked the following: the wild-type one-letter amino acid code (amino_acid_clean), missense pairs (WT;Mut) with the lowest cost (min_cost_mutant) and missense pairs with the highest cost (max_cost_mutant).

*Physicochemical features:* Amino acid identity was extracted from the Missense csv file and stored as the one-letter code for each position ($i$). All amino acid identities were encoded as numerical values (1 or 0) in distinct columns for each unique value (e.g. 20 columns). Amino acid charges were hard coded where D, E= –1, K, R, H= +1, and all others were 0. Hydrophobicity was defined at every position using the amino acid identity, where hydropathy indexes were I: 4.5, V: 4.2, L: 3.8, F: 2.8, C: 2.5, M: 1.9, A: 1.8, G: -0.4, T: -0.7, S: -0.8, W: -0.9, Y: -1.3, P: -1.6, H: -3.2, E: -3.5, Q: -3.5, D: -3.5, N: -3.5, K: -3.9, and R: -4.5 (*72*). Catalytic capacity was also flagged, where D, E, C, Y, H= can_act_as_acid and H, K, R, C, Y= can_act_as_base.

*z-scores for key attributes:* To facilitate downstream position ($i$) ranking, we calculated a z-score for select attributes using the standard equation:

$$z_i(x) = \frac{x_i - \mu_x}{Var_x}$$

where $x$= RSA, WCN, or effect score and returning z-scores were defined as rsa_z, wcn_z, and effect_z, respectfully, for each residue of a protein.

*Linear sequence features:* We defined a linear sequence window ($W_i$) where for each position ($i$) we included up to five amino acids (excluding itself) comprising positions both up and downstream as defined by $W_i = \{j \mid |j - i| \leq 5, j \neq i\}$. Applying this sequence window, we calculated:

i) how many significant positions were identified within the sequence window of position $i$

$$\text{Seq\_n\_sig\_w5} = \sum_{j \in w_i} sig_j$$

ii) the fraction of significant positions within the sequence window of position $i$

$$\text{Seq\_frac\_sig\_w5} = \frac{\text{seq\_n\_sig\_w5}_{\text{i}}}{|w_i|}$$

iii) the mean effect score of all positions within the sequence window of position $i$

$$\text{Seq\_mean\_effect\_w5} = \frac{1}{|w_i|} \sum_{j \in w_i} effect_j$$

*Three-dimensional context features:* We defined the three-dimensional neighborhood of each position ($i$) by:

$$N_i = \{j \mid \left|\left|r_j - r_i\right|\right| \mid \leq 9\text{Å}, j \neq i\}$$

where $N_i$= set of amino acids whose C-alpha carbon is less than 9 Å from the C-alpha carbon of position $i$, $r_i$= the coordinates (x, y, z) of the C-alpha carbon of position $i$, and $r_j$= the coordinates (x, y, z) of the C-alpha carbon of position $j$. We then computed six three-dimensional derived attributes using the following equations:

1) total number of 3D neighbors: $\text{n\_3D\_neighbors}_i = |N_i|$
2) total number of significant 3D neighbors: $\text{n\_3D\_sig\_neighbors}_i = \sum_{j \in N_i} sig_j$
3) fraction of significant 3D neighbors: $\text{frac\_3D\_sig\_neighbors}_i = \frac{\text{n\_3D\_sig\_neighbors}_i}{n\_3D\_neighbors_i}$
4) mean effect score of 3D neighbors: $\text{mean\_3D\_effect\_neighbors}_i = \frac{1}{|N_i|} \sum_{j \in w_i} effect_j$
5) median effect score of 3D neighbors: $\text{median\_3D\_effect\_neighbors}_i = \text{median}\left(\{effect_j \mid j \in N_i\}\right)$
6) maximum effect score of 3D neighbors: $\text{max\_3D\_effect\_neighbors}_i = \max\left(\{effect_j \mid j \in N_i\}\right)$

*Log transform residue features:* To enhance performance of downstream machine-learning, we log transformed the following position ($i$) metrics:

1) p-value: $\text{neglog}10\_p\_value_i = -\log_{10} p_i$
2) q-value: $\text{neglog}10\_q\_value_i = -\log_{10} q_i$
3) min_cost: $\text{min\_cost\_log1p}_i = \log(1 + \text{min\_cost}_i)$
4) max_cost: $\text{max\_cost\_log1p}_i = \log(1 + \text{max\_cost}_i)$
5) rsa: $\text{rsa\_log1p}_i = \log(1 + \text{rsa}_i)$
6) pae: $\text{pae\_log1p}_i = \log(1 + \text{pae}_i)$

7) total number of 3D neighbors: $\text{n\_3d\_neightbors\_log1p}_i = \log(1 + \text{n\_3d\_neightbors}_i)$

8) total number of significant 3D neighbors: $\text{n\_3D\_sig\_neighbors\_log1p}_i = \log(1 + \text{n\_3D\_sig\_neighbors}_i)$

9) total significant positions within the sequence window: $\text{seq\_n\_sig\_w5\_log1p}_i = \log(1 + \text{seq\_n\_sig\_w5}_i)$

Residue-level metrics and statistics can also be viewed using a volcano plot where $Effect_i$ is defined on the x-axis (x) and a clamped p-value ($p_{clamped_i}$) is shown on the y-axis (y). The $p_{clamped_i}$ value is defined for each residue as:

$$p_{clamped_i} = \{p_i \text{ } if \text{ } p_i > 0 \text{ } and \text{ } not \text{ } NAN \mid \min\{p_j \mid p_j > 0\} \text{ } if \text{ } p_i \le 0 \text{ } or \text{ } NaN\}$$

where NAN= not a number, $p_i$= p-value of residue $i$, and $p_j$= p-value of residue j.

Model Training

The TAMALE model was trained on a curated subset of human proteins. The UniProt database listed 2,307 reviewed human proteins with one or more annotated active site residues (*10*). From this initial set, 2,264 also had AlphaMissense data available through the AlphaFold database (*1*). The revised list was further divided into training data comprised of 1,356 human proteins, validation data totaling 453, and test data of 453 targets (**data S1**). The split strategy for creating these representative sets was to balance enzyme commission number, protein length, and AlphaFold confidence. Using the assembled residue feature table and the reviewed UniProt active site residue annotations as ground truth, we trained a light gradient boosting model (LGBM) utilizing an open-source implementation from the Scikit-learn python package (*11, 73*). For each residue position in the target protein, the model outputs and appends a newly created TAMALE score in the feature table. TAMALE scores are used to rank each residue by predicted relative importance. Both the raw TAMALE score and TAMALE-based ranking are utilized in downstream analysis for characterization and discovery across the human proteome.

TAMALE Benchmarking

The test set was never used during model training and comprises 453 reviewed human proteins with one or more annotated active site residue (data S1). Benchmarking metrics reporting the performance of the TAMALE model used the test set, demonstrating the usefulness of both the raw TAMALE score for discriminating annotated active site residues from non-annotated residues and the TAMALE-based ranking for identify residues of functional importance. To assess model stability, the training and validation sets were combined to generate 100 randomized subsets for model training and benchmarking. TAMALE was further benchmarked across all seven general enzyme classes, AlphaFold model confidence, and protein length, ranging from typical polypeptides (400-500 aa) to long (>2,000 aa) polypeptides. Performance was also assessed for the identification of active site residues, including catalytic residues, as well as metal and ligand binding residues, and regulatory and allosteric residues. Compute time for TAMALE analyses was calculated using a standard MacBook Pro laptop with an M5 chip and 16GB memory and for EasiFA using its GPU-supported server at http://cadd.iddd.group/easifa/ (*6*).

FASTKD5 Expression and Purification

Recombinant human FASTKD5 encodes residues 111-764 to resemble the mature, proteolytically processed mitochondrial matrix protein. Sequence verified plasmid DNA harbored insect cell codon optimized FASTKD5, cloned in frame with an N-terminal $His_6$ affinity tag followed by a TEV protease recognition site in the pFasBacHT-B vector (GenScript). Please refer to table S2 for a list of all expression plasmids used in this study. Recombinant baculovirus production and FASTKD5 variant expression were performed using *Spodoptera frugiperda* Sf9 cells (Gibco, catalog 11496015, lot 3138016) following manufacturer guidelines. Insect cells were maintained at 27 ºC, 125 rpm and harvested 3 days post-infection (Invitrogen). Infected insect cells were resuspended in 50 mM Tris pH 7.4, 500 mM NaCl, 5 mM 2-mercaptoethanol, 15 mM imidazole, 2% (v/v) glycerol, 1 mM benzamidine, 1 mM phenylmethylsulfonyl fluoride, 5 μg/mL leupeptin, and 0.7 μg/mL pepstatin A. Resuspended cells were lysed by sonication at 40% amplitude for 1 minute (5 second pulses) at 4 °C. Lysate was clarified at 8,000 x g for 30 minutes at 4 °C. A two-step purification protocol, consisting of affinity chromatography followed by size exclusion chromatograph, resulted in purified recombinant FASTKD5 protein. Clarified lysate was incubated with His60 nickel resin (Takara) for 30 minutes at 4 °C. The resin was washed five times with ten column volume of 50 mM Tris pH 7.4, 500 mM NaCl, 5 mM 2-mercaptoethanol, and 30 mM imidazole. $His_6$-tagged FASTKD5 protein was eluted in 50 mM Tris pH 7.4, 500 mM NaCl, 5 mM 2-mercaptoethanol, 300 mM imidazole, and 2% (v/v) glycerol and resolved over a Superdex 200 HiLoad 16/600 column equilibrated with 50 mM Tris pH 7.4, 500 mM NaCl, 5 mM 2-mercaptoethanol, and 2% (v/v) glycerol. Purified FASTKD5 variants were flash frozen using liquid nitrogen and stored at -80 °C.

*In vitro* FASTKD5 RNA Cleavage Assay

FASTKD5 RNA cleavage activity was measured *in vitro* using a model RNA substrate of the ATP8/6-CO3 mitochondrial mRNA junction alongside a custom 3′-fluorescein labeled RNA ladder. Please refer to table S3 for RNA sequences. The synthetic substrate encodes a 5′-Cy5 fluorophore and 3′-fluorescein fluorophore for RNA visualization and fragment discrimination (Horizon Discovery Biosciences). ATP8/6-CO3 RNA (100 nM) was mixed with FASTKD5 variants (50 nM) in reaction buffer (50 mM Tris pH 7.5, 100 mM NaCl, 5 mM 2-mercaptoethanol, and 5 mM EDTA) and incubated for 5 minutes at 37 °C. The reaction was quenched by supplementing the mixture with equal volume of urea loading dye and 1.6 units of proteinase K and incubated for 15 minutes at 25 °C. Samples were heated to 75 °C for 15 minutes and resolved using a 15% urea polyacrylamide gel. Intact RNA substrate and cleavage products were visualized using a ChemiDoc MP imaging system (BioRad). Three independent technical replicates were performed for each RNA cleavage reaction condition.

*In vitro* FASTKD5 RNA Binding Assay

Biolayer interferometry (BLI) was performed at 27 °C using an Octet R8 instrument and Streptavidin (SA) biosensor (Sartorius). A 3′-biotinylated RNA substrate (Horizon Discovery Biosciences) comprising the sequence spanning the ATP8/6-CO3 mitochondrial mRNA junction was used to measure FASTKD5 RNA binding activity *in vitro*. Please refer to table S3 for the

RNA sequence. Biotinylated RNA (100 nM) and a series of FASTKD5 variants diluted to a final concentration of 35.6, 71.3, 143, 285, and 570 nM were prepared using RNA binding buffer (50 mM Tris pH 7.5, 50 mM NaCl, 5 mM 2-mercaptoethanol, 0.05% (v/v) Tween 20). Prior to the experiment, biosensors were equilibrated in 200 μL of RNA binding buffer for 10 minutes. The BLI experiment comprised the following steps using 200 μL sample volumes: biosensor initial baseline in RNA binding buffer (60 seconds), substrate loading in 100 nM biotinylated RNA (300 seconds), secondary baseline in RNA binding buffer (30 seconds), FASTKD5 association in 35.6-570 nM protein dilutions (300 seconds), and FASTKD5 dissociation in RNA binding buffer (300 seconds). Baseline subtraction and data fitting using the Langmuir 1:1 kinetics model was performed with the Octet Analysis Studio version 12.2.2.26 software (Sartorius). Savitzky-Golay filtering was applied. Two independent technical replicates were performed, and the mean and standard deviation are reported in Fig. 5.

Mass Photometry

The molecular mass of recombinant FASTKD5 was determined using a TwoMP mass photometer (Refeyn). Glass coverslips were prepared as previously described (*74*). A FASTKD5 protein mixture was prepared using 50 mM Tris pH 7.4, 100 mM NaCl, and 5 mM 2-mercaptoethanol to a final concentration of 25 nM. The diluted sample was spun at 21,300 x g for 10 minutes immediately prior to the mass photometry measurement. Acquire MP software was used for acquiring a 60 second recording with a total count ranging between 500-2,000. A MassFerence P1 calibrant (Refeyn) was analyzed and applied to the FASTKD5 data using DiscoverMP software to calculate a molecular mass. A representative plot is illustrated in fig. S15 from three independent technical replicates.

**Supplementary Text**

Analysis of USP34

We evaluated TAMALE on an extreme case, the 3,546-residue USP34 deubiquitinating enzyme. Ubiquitin-specific proteases (USPs) regulate protein degradation by removing ubiquitin from substrates. TAMALE correctly ranked the catalytic residue H2164 as the top prediction and C1903 as the second, consistent with their proposed roles in the catalytic dyad and structural rearrangement upon ubiquitin binding (**fig. S9**) (*75*). While N2187 was moderately ranked (33$^{rd}$), a nearby residue, D2188 (4$^{th}$), occupied a comparable position relative to the catalytic histidine and corresponds to a conserved functional contact observed in other USPs, suggesting a potential role in stabilizing the transition state (*76, 77*). The model also recovered known ubiquitin binding residues, such as D1978, E1981, and Y2165 (*75*). Collectively, this analysis demonstrates the strong performance of TAMALE on very large proteins, accurately identifying catalytic and substrate binding residues.

Analysis of ADRB2

We tested ADRB2, a prototypical GPCR, and allosteric signaling receptor (*78*). Twenty of the top 25 residues cluster within known regulatory regions, including the orthosteric binding site ($D^{3.32}$ (2$^{nd}$), $N^{7.39}$ (14$^{th}$), $N^{6.55}$ (17$^{th}$)), conserved features for agonist ligand binding ($P^{5.50}$ (31$^{st}$), $S^{5.46}$ (16$^{th}$)), key allosteric elements such as the sodium ion binding site and conserved water network ($D^{2.50}$ (1$^{st}$), $S^{3.39}$ (3$^{rd}$)), and microswitches for G-protein binding. This analysis demonstrates TAMALE's capacity to identify regulatory sites in non-catalytic proteins.

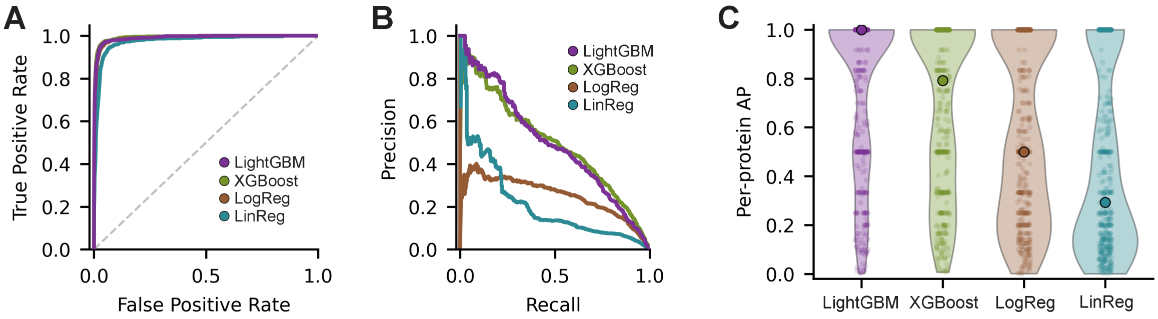


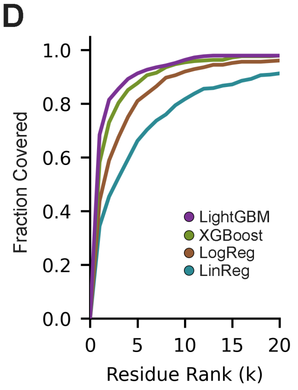


E

| | | | | Protein Recovery | | | | Residue Recovery | | | |
|---|---|---|---|---|---|---|---|---|---|---|---|
| Model | AUROC | AUPRC | AP | K=1 | K=5 | K=10 | K=20 | K=1 | K=5 | K=10 | K=20 |
| LightGBM | 0.991 | **0.494** | **0.753** | **0.685** | **0.913** | **0.963** | **0.979** | **0.454** | **0.858** | **0.939** | **0.970** |
| XGBoost | **0.993** | **0.494** | 0.670 | 0.582 | 0.877 | 0.954 | **0.979** | 0.386 | 0.806 | 0.923 | 0.968 |
| LogReg | 0.988 | 0.252 | 0.561 | 0.438 | 0.811 | 0.920 | 0.961 | 0.290 | 0.714 | 0.884 | 0.944 |
| LinReg | 0.975 | 0.209 | 0.435 | 0.345 | 0.662 | 0.817 | 0.913 | 0.228 | 0.531 | 0.711 | 0.871 |

**fig. S1. Supervised machine learning model performance in active site prediction.** (**A**) Receiver operating characteristic curve (ROC) for Light Gradient Boosting Machine (purple, LightGBM), eXtreme Gradient Boosting (XGBoost, green), Logistic Regression (LogReg, brown), and Linear Regression (LinReg, cyan). Dotted line is the baseline for random chance. (**B**) Precision-recall curve (PRC) for predicting all annotated active site residues within a given target protein. (**C**) Average precision (AP) for predicting a protein that contains an active site residue. (**D**) Recovery curve of active site amino acids by ranked residue position. (**E**) Performance metrics of active site prediction across models where K represents the top ranked residues.

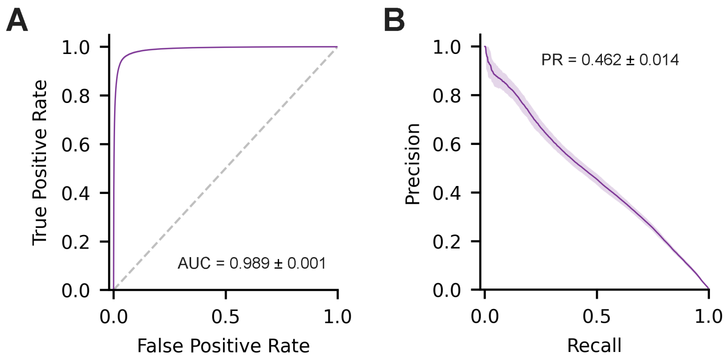


**fig. S2. LightGBM model performance across resampling.** (**A**) Mean receiver operating characteristic curve (ROC) area under the curve (AUC). (**B**) Mean precision-recall curve. Mean and variance were calculated from 100 iterations of model training from randomized subsets.

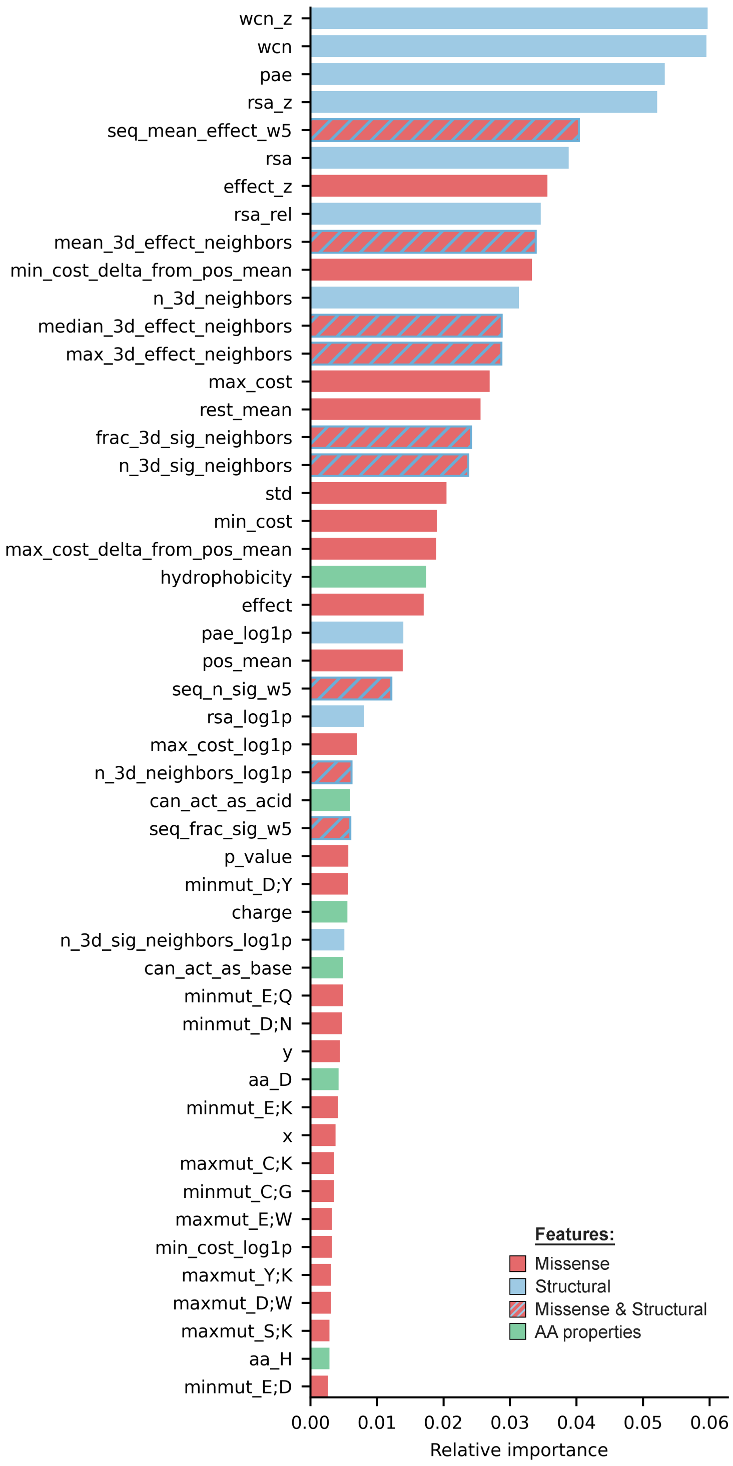


**fig. S3. Relative importance of top 50 residue features in the TAMALE model.** Residue attributes and statistics driving performance of the TAMALE model.

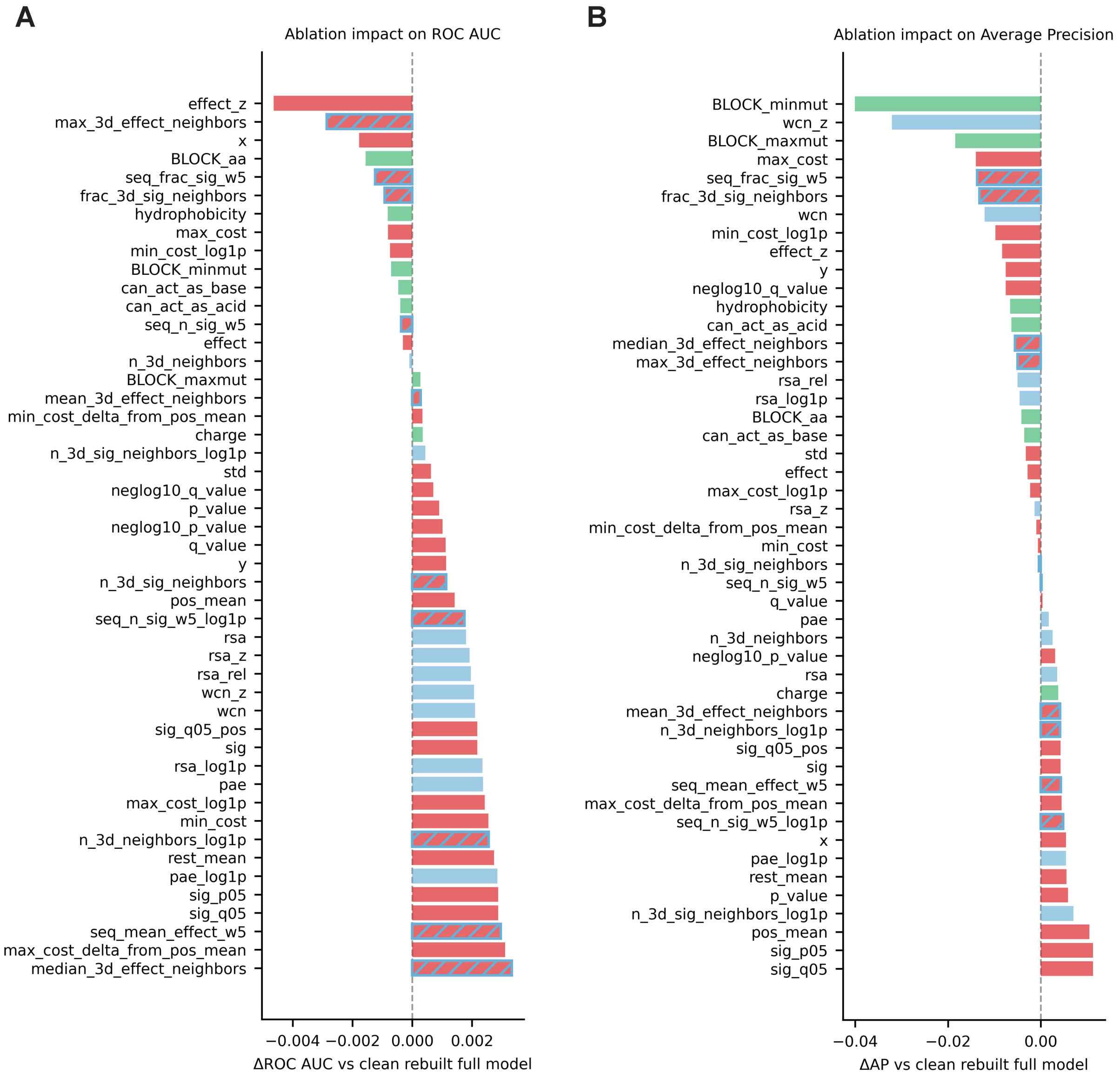


**fig. S4. Ablation study on the role of individual residue-level features.** Comparison of ROC and AP between TAMALE and model variants lacking an individual residue-level feature.

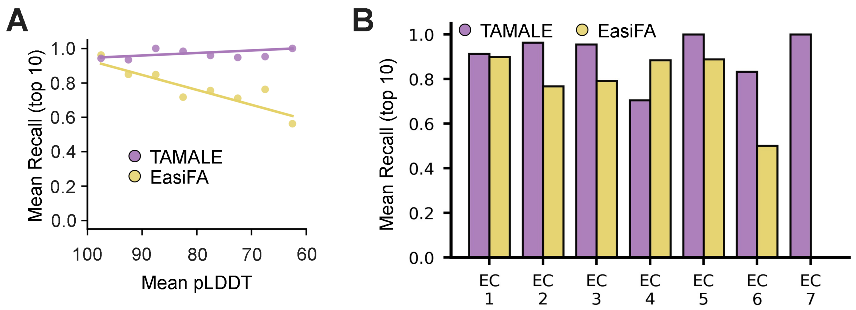


**fig. S5. Performance metrics of TAMALE and EasiFA.** (**A**) Recall using the top 10 positions for TAMALE and EC-guided EasiFA relative to AlphaFold model prediction confidence. (**B**) Protein-level identification of annotated active site residues within the top 10 positions based on enzyme commission number (EC) where EC 1 is oxidoreductases, EC 2 is transferases, EC 3 is hydrolases, EC 4 is lyases, EC 5 is isomerases, EC 6 is ligases, and EC 7 is translocases.

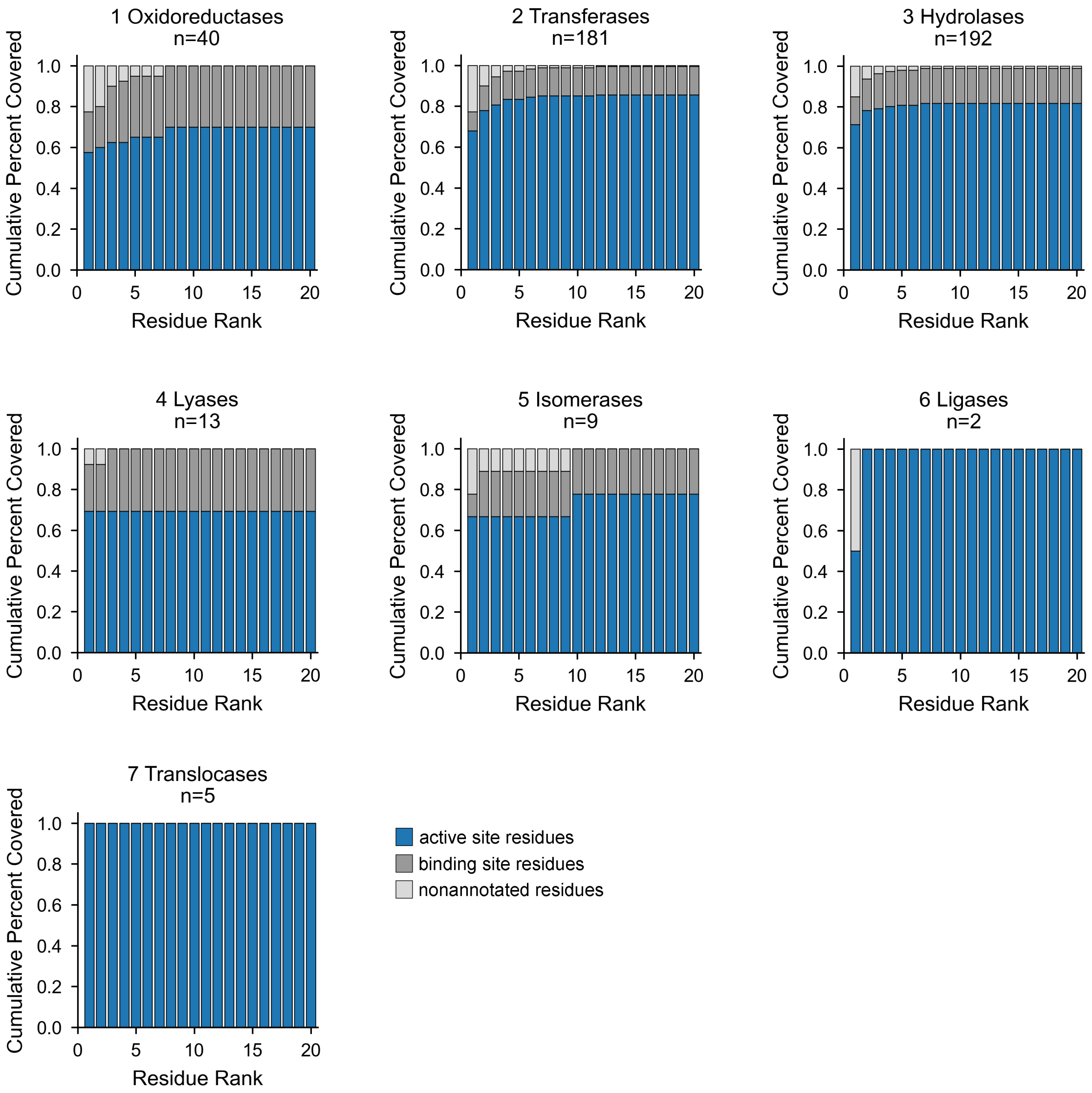


**fig. S6. TAMALE performance with annotated binding residues.** Enzyme class specific plots depicting identification of annotated binding residues (dark grey) and active site residues (blue) across residue rank. EC 1 is oxidoreductases, EC 2 is transferases, EC 3 is hydrolases, EC 4 is lyases, EC 5 is isomerases, EC 6 is ligases, and EC 7 is translocases.

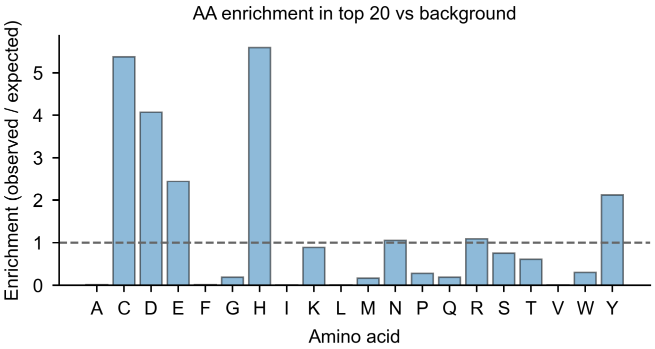


**fig. S7. Amino acid enrichment by TAMALE.** Plot depicting relative enrichment of each amino acid across the top 20 TAMALE-ranked residue positions. The dotted line represents the baseline expectancy in the absence of enrichment or deprioritization.

**A**

| Enzyme Class | Target | Active Site | TAMALE Ranking | TAMALE Score |
|---|---|---|---|---|
| EC 1 :oxidoreductase | D-3-phosphoglycerate dehydrogenase | H283 | 1st of 533 | 2.88 |
| | | H265 | 4th of 533 | 0.59 |
| | | R236 | 9th of 533 | −0.71 |
| EC 2: transferase | serine/threonine-protein kinase haspin | D649 | 1st of 798 | 5.51 |
| EC 3: hydrolase | protein-arginine deiminase type-4 | C645 | 1st of 663 | 4.48 |
| | | D350 | 2nd of 663 | 3.03 |
| | | D473 | 5th of 663 | 1.64 |
| | | H471 | 6th of 663 | 1.54 |
| EC 4: lyase | S-adenosylmethionine decarboxylase proenzyme | E11 | 1st of 334 | 3.08 |
| | | H243 | 2nd of 334 | 2.94 |
| | | C82 | 3rd of 334 | 1.62 |
| | | S68 | 9th of 334 | −1.41 |
| | | E8 | 13th of 334 | −2.48 |
| | | S229 | 17th of 334 | −2.94 |
| EC 5: isomerase | DNA topoisomerase 2-beta | D562* | 1st of 1626 | 2.53 |
| | | Y826 | 2nd of 1626 | 1.90 |
| | | D564* | 9th of 1626 | −0.34 |
| EC 6: ligase | GMP synthase | C104 | 1st of 693 | 3.51 |
| | | H190 | 5th of 693 | −0.75 |
| | | E192 | 36th of 693 | −4.00 |
| | DNA ligase I | D570* | 1st of 919 | 2.81 |
| | | E621* | 2nd of 919 | 1.80 |
| | | K568 | 3rd of 919 | 1.74 |
| EC 7: translocase | probable phospholipid-transporting ATPase IIA | D391 | 1st of 1047 | 1.98 |

**B** **DUSP28 (Phosphatase)**

| Rank | Residue | Score | Annot. | Function |
|---|---|---|---|---|
| 1 | D72 | 0.66 | N | Active Site |
| 2 | R109 | 0.16 | N | Active Site |
| 3 | S108 | -0.75 | N | Unknown |
| 4 | C103 | -0.81 | Y | Active Site |
| 5 | S110 | -2.61 | N | Unknown |

**E** **MRE11 (Exonuclease)**

| Rank | Residue | Score | Annot. | Function |
|---|---|---|---|---|
| 1 | H245 | 3.77 | Y | Binding |
| 2 | H129 | 3.63 | Y | Active Site |
| 3 | H217 | 3.09 | Y | Binding |
| 4 | H22 | 2.70 | Y | Binding |
| 5 | H247 | 2.53 | Y | Binding |

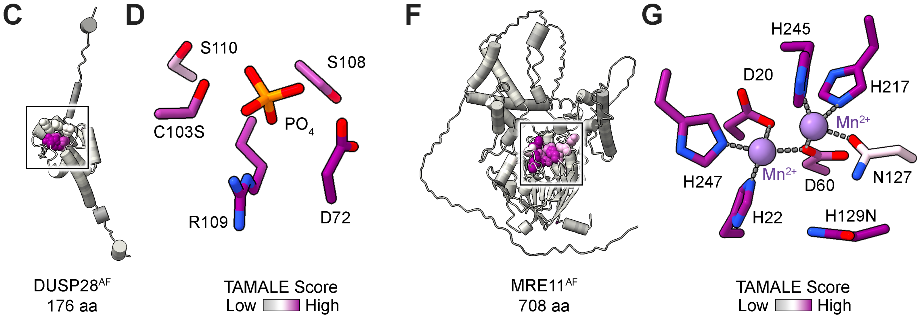


**fig. S8. Case studies of representative enzymes across enzyme class.** (**A**) Summary of TAMALE rankings for annotated active site residues in representative enzymes across enzyme class. Asterisk marks active site residues that lack annotation in the UniProt database. (**B**) Top 5 TAMALE-predicted DUSP28 functional residues (UniProt ID Q4G0W2). Annotated active site residues (Annot.) in the UniProt database are defined by Y for yes and missing active site annotations are defined by N for no. (**C**) DUSP28 AlphaFold (AF) model prediction with TAMALE coloring reveals a cluster demarcated by a box (*1*). (**D**) Phosphate-bound DUSP28 catalytic site with TAMALE coloring (PDB ID 5Y15; (*19*)). (**E**) Top 5 TAMALE-predicted MRE11 functional residues (UniProt ID P49959). Annotated binding residues and active site residues in the UniProt database are defined by Y for yes. (**F**) MRE11 AF model prediction with TAMALE coloring reveals a cluster demarcated by a box (*1*). (**G**) Manganese-bound MRE11 catalytic site with TAMALE coloring (PDB ID 8BAH; (*20*)).

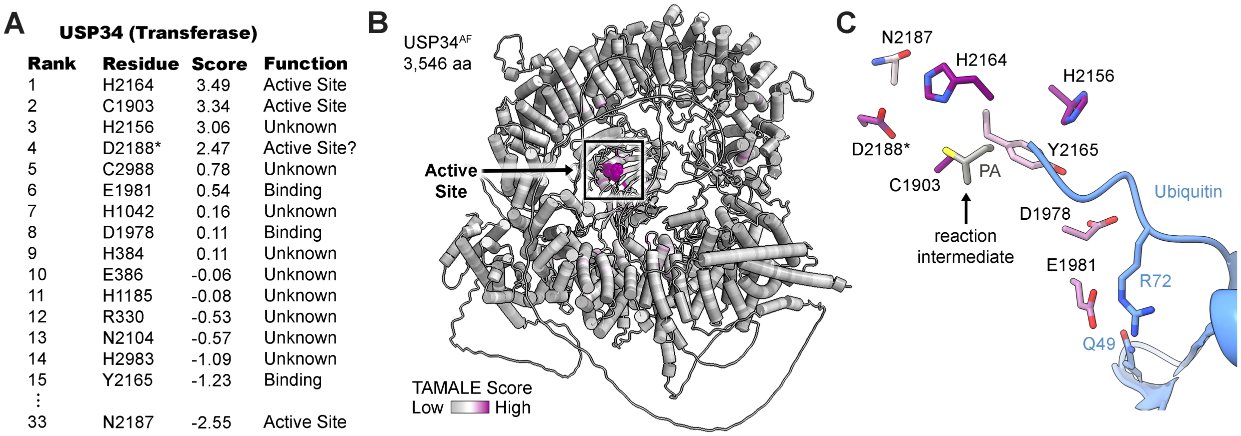


**fig. S9. Active site identification in the 3,546 aa USP34 enzyme.** (**A**) Top TAMALE-predicted USP34 functional residues (UniProt ID Q70CQ2). (**B**) USP34 AlphaFold (AF) model prediction with TAMALE coloring reveals a discrete cluster demarcated by a box (*1*). (**C**) Ubiquitin substrate-bound USP34 with TAMALE coloring. Ubiquitin is shown in blue (PDB ID 7W3U, (*75*)) and propargylamine (PA) was covalently linked to the catalytic cysteine forming a vinyl thioester meant to mimic a reaction intermediate (*75*).

**A** **UBE2D3 (ubiquitin ligase)**

| Rank | Residue | Score | Function |
|---|---|---|---|
| 1 | C85 | 0.73 | Active Site |
| 2 | D87 | -2.43 | Active Site |
| 3 | D112 | -3.31 | Unknown |
| 4 | D117 | -4.20 | Active Site |
| 5 | R5 | -4.45 | Binding |
| 6 | C21 | -4.46 | Unknown |
| 7 | E9 | -4.66 | Binding |
| 8 | S94 | -5.07 | Unknown |
| 9 | C107 | -5.14 | Unknown |
| 10 | H75 | -5.17 | Unknown |

**D** **BRCA1 (ubiquitin ligase scaffold)**

| Rank | Residue | Score | Function |
|---|---|---|---|
| 1 | C44 | 3.30 | Binding |
| 2 | C47 | 2.18 | Binding |
| 3 | D1840 | 2.12 | Regulatory |
| 4 | C24 | 1.99 | Binding |
| 5 | C1697 | 1.87 | Regulatory |
| 6 | C27 | 1.41 | Binding |
| 7 | H41 | 1.00 | Binding |
| 8 | D1739 | 0.83 | Unknown |
| 9 | C1768 | 0.49 | Unknown |
| 10 | E1735 | 0.45 | Unknown |
| 11 | H1746 | 0.42 | Unknown |
| 12 | E1836 | -0.32 | Regulatory |
| 13 | C64 | -0.51 | Binding |
| 14 | C39 | -0.52 | Binding |
| 15 | C61 | -0.60 | Binding |

**H** **BARD1 (ubiquiting ligase scaffold)**

| Rank | Residue | Score | Function |
|---|---|---|---|
| 1 | C733 | 1.55 | Unknown |
| 2 | D458 | 0.70 | Binding |
| 3 | E467 | 0.40 | Binding |
| 4 | H723 | -0.13 | Unknown |
| 5 | H606 | -0.29 | Unknown |
| 6 | D500 | -0.52 | Binding |
| 7 | C675 | -0.59 | Unknown |
| 8 | H68 | -1.06 | Binding |
| 9 | E429 | -1.31 | Binding |
| 10 | H473 | -1.39 | Unknown |
| 11 | D495 | -1.41 | Binding |
| 12 | H433 | -1.63 | Binding |
| 13 | D673 | -1.88 | Binding |
| 14 | H506 | -2.32 | Unknown |
| 15 | H466 | -2.44 | Binding |

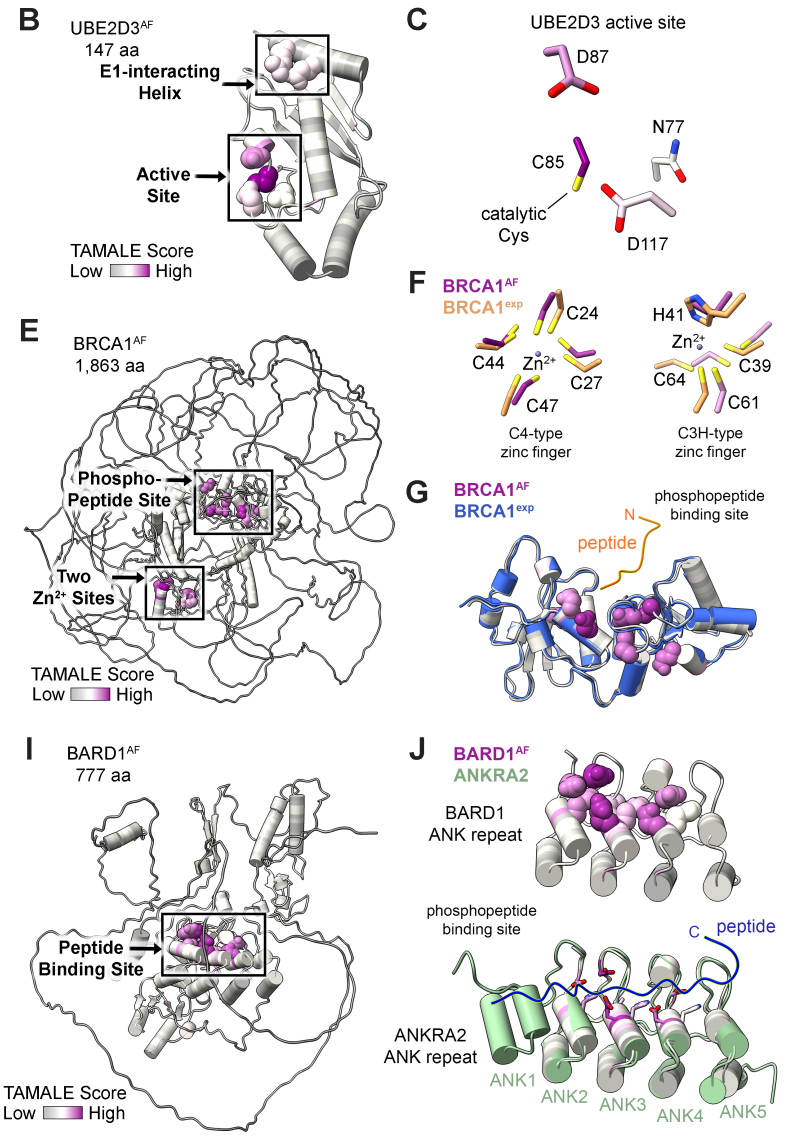


**fig. S10. Functional site detection of molecular scaffolds.** (**A**) Top 10 TAMALE-predicted UBE2D3 functional residues (UniProt ID P61077). (**B**) UBE2D3 AlphaFold (AF) model prediction with TAMALE coloring reveals a cluster demarcated by a box (*1*). (**C**) Active site of the UBE2D3 E2 ubiquitin-conjugating enzyme in TAMALE color (PDB ID 7LYB; (*24*)). (**D**) Top 15 TAMALE-predicted BRCA1 functional residues (UniProt ID P38398). (**E**) BRCA1 AlphaFold (AF) model prediction with TAMALE coloring reveals three discrete clusters demarcated by two boxes (*1*). (**F-G**) Two zinc ion sites and a phosphopeptide binding site of BRCA1 in TAMALE color and overlay of experimental (exp) data (PDB ID 4U4A, 1JM7; (*79*)). (**H**) Top 15 TAMALE-predicted BARD1 functional residues (UniProt ID Q99728). (**I**) BARD1 AlphaFold (AF) model prediction with TAMALE coloring reveals a prominent cluster in the ankyrin (ANK) repeat (*1*). (**J**) Overlay of the BARD1 ANK repeat shown in TAMALE colors with the peptide-bound ANKRA2 ANK repeat (PDB ID 7LYB, 3V31; (*24, 80*)).

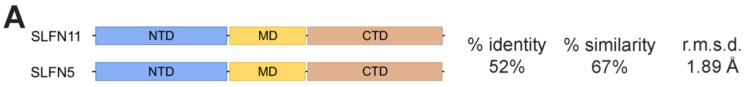


**B SLFN11 (Endoribonuclease)**

| Rank | Residue | Score | Function |
|---|---|---|---|
| 1 | K216 | 2.51 | Active Site |
| 2 | E669 | 0.89 | Binding |
| 3 | E209 | 0.78 | Active Site |
| 4 | H664 | -0.33 | Unknown |
| 5 | K605 | -0.38 | Binding |
| 6 | E861 | -0.61 | Unknown |
| 7 | E214 | -0.67 | Active Site |
| 8 | D703 | -0.76 | Unknown |
| 9 | E632 | -0.81 | Unknown |
| 10 | D668 | -0.88 | Binding |
| 11 | C51 | -0.94 | Unknown |
| 12 | C884 | -0.98 | Unknown |
| 13 | C321 | -1.03 | Binding |
| 14 | H598 | -1.13 | Unknown |
| 15 | T606 | -1.83 | Binding |

**C SLFN5 (Pseudo-endoribonuclease)**

| Rank | Residue | Score | Function |
|---|---|---|---|
| 1 | S878 | 0.22 | Unknown |
| 2 | C302 | 0.12 | Binding |
| 3 | H645 | -0.01 | Unknown |
| 4 | C48 | -0.45 | Unknown |
| 5 | D789 | -0.73 | Unknown |
| 6 | E852 | -0.83 | Unknown |
| 7 | D684 | -1.03 | Unknown |
| 8 | H266 | -1.12 | Binding |
| 9 | D428 | -1.54 | Unknown |
| 10 | S422 | -1.56 | Unknown |
| 11 | C610 | -1.67 | Unknown |
| 12 | D649 | -1.75 | Binding? |
| ⋮ | | | |
| 16 | T585 | -1.97 | Binding? |

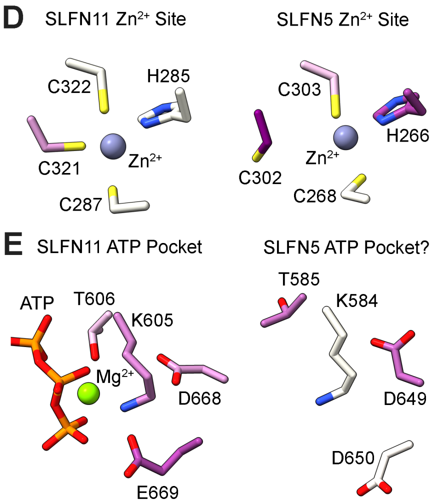


**fig. S11. TAMALE analysis of SLFN proteins identify a structural ion site and ATP binding pocket.** (**A**) Illustration of domain organization of SLFN proteins SLFN11 and SLFN5 where the N-terminal domain (NTD) is blue, the Middle domain (MD) is yellow, and the C-terminal domain (CTD) is brown. The human SLFN11 and SLFN5 protein sequences were analyzed by BLAST to calculate sequence identity and similarity (*2*). The root mean square deviation (r.m.s.d.) of experimental structures were calculated using monomeric SLFN11 (PDB ID 9ERD; (*30*)) and SLFN5 (PDB ID 7PPJ; (*29*)). (**B**) Top 15 TAMALE-predicted SLFN11 endoribonuclease functional residues (UniProt ID Q7Z7L1). (**C**) TAMALE-predicted SLFN5 pseudo-endoribonuclease functional residues (UniProt ID Q08AF3). (**D**) Validated $Zn^{2+}$ metal binding sites in SLFN11 and SLFN5 (*28, 29*). (**E**) Validated SLFN11 ATP-bound pocket with consensus P-loop and Walker B motif residues alongside the putative SLFN5 ATP pocket (*30*).

**A** **KRAS (GTP Hydrolase)**

| Rank | Residue | Score | Function |
|---|---|---|---|
| 1 | D57 | 1.71 | Active Site |
| 2 | K16 | -0.56 | Active Site |
| 3 | D38 | -0.84 | Binding |
| 4 | D54 | -1.56 | Regulatory |
| 5 | D33 | -1.61 | Binding |
| 6 | T58 | -2.18 | Regulatory |
| 7 | K117 | -2.57 | Binding |
| 8 | D69 | -3.28 | Regulatory |
| 9 | E37 | -3.56 | Binding |
| 10 | Y32 | -3.67 | Regulatory |
| 11 | E63 | -3.75 | Binding |
| 12 | D119 | -3.87 | Binding |
| 13 | E143 | -4.03 | Unknown |
| 14 | N116 | -4.43 | Binding |
| 15 | Y71 | -4.45 | Regulatory |
| 16 | G15 | -4.65 | Active Site |
| 17 | E62 | -4.66 | Regulatory |
| 18 | T35 | -4.77 | Regulatory |
| 19 | R68 | -4.90 | Active Site |
| 20 | G60 | -5.19 | Regulatory |
| 21 | S17 | -5.29 | Active Site |
| 22 | Y40 | -5.33 | Regulatory |
| 23 | D47 | -5.35 | Regulatory |
| 24 | E3 | -5.63 | Regulatory |
| 25 | S39 | -5.74 | Regulatory |

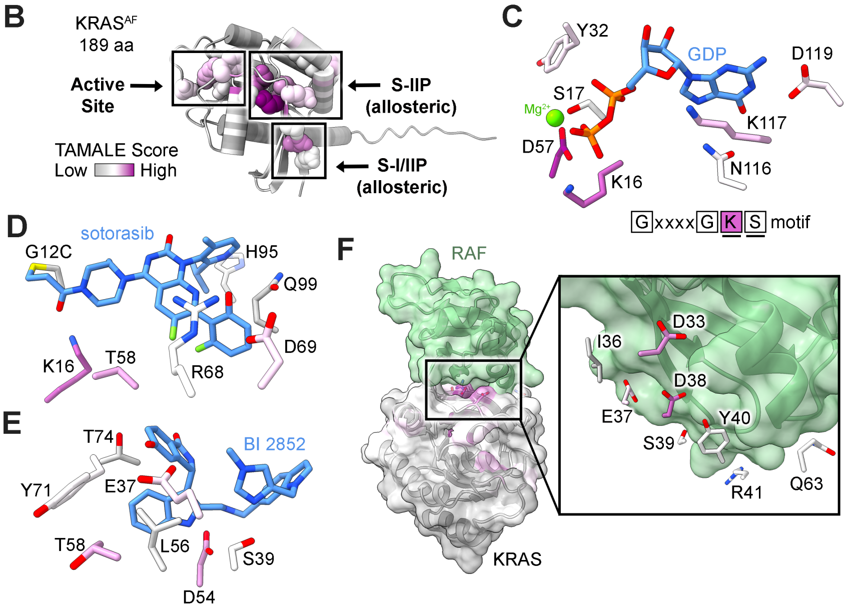


**fig. S12. TAMALE correctly predicts regulatory sites in KRAS.** (**A**) Top 25 TAMALE-predicted KRAS functional residues (UniProt ID P01116). (**B**) KRAS AlphaFold (AF) model prediction with TAMALE coloring demonstrates three distinct clusters demarcated by boxes and representing the nucleotide binding pocket and two druggable allosteric pockets (*1*). (**C**) KRAS nucleotide pocket and P-loop (GxxxGKS) motif colored by TAMALE score (PDB ID 6GJ8; (*33*)). (**D-E**) Druggable KRAS allosteric pockets bound to FDA-approved sotorasib (blue, PDB ID 6OIM; (*81*)) and inhibitor, BI 2825 (blue, PDB ID 6GJ8; (*33*)). (**F**) KRAS-effector binding interface where the RAF proto-oncogene is colored in green (PDB ID 6VJJ; (*36*)).

**A** **ADRB2 (Cell surface receptor)**

| Rank | Residue | BW | Score | Function |
|---|---|---|---|---|
| 1 | D79 | $D^{2.50}$ | 1.03 | Regulatory |
| 2 | D113 | $D^{3.32}$ | 0.82 | Regulatory |
| 3 | S120 | $S^{3.39}$ | -1.04 | Regulatory |
| 4 | C285 | $C^{6.47}$ | -1.43 | Regulatory |
| 5 | S319 | $S^{7.46}$ | -1.46 | Regulatory |
| 6 | R131 | $R^{3.50}$ | -1.49 | Regulatory |
| 7 | N51 | $N^{1.50}$ | -1.62 | Regulatory |
| 8 | Y219 | $Y^{5.58}$ | -1.82 | Regulatory |
| 9 | Y141 | $Y^{34.53}$ | -2.28 | Unknown |
| 10 | H269 | $H^{6.31}$ | -2.53 | Unknown |
| 11 | C116 | $C^{3.35}$ | -2.62 | Regulatory |
| 12 | D130 | $D^{3.49}$ | -2.76 | Regulatory |
| 13 | C106 | $C^{3.25}$ | -3.20 | Unknown |
| 14 | N312 | $N^{7.39}$ | -3.49 | Regulatory |
| 15 | P323 | $P^{7.50}$ | -3.52 | Unknown |
| 16 | S207 | $S^{5.46}$ | -3.63 | Regulatory |
| 17 | N293 | $N^{6.55}$ | -3.70 | Regulatory |
| 18 | Y326 | $Y^{7.53}$ | -3.87 | Regulatory |
| 19 | S329 | $S^{8.47}$ | -3.97 | Unknown |
| 20 | N322 | $N^{7.49}$ | -4.00 | Unknown |
| 21 | C125 | $C^{3.44}$ | -4.15 | Regulatory |
| 22 | T118 | $T^{3.37}$ | -4.31 | Regulatory |
| 23 | N318 | $N^{7.45}$ | -4.43 | Unknown |
| 24 | F282 | $F^{6.44}$ | -4.51 | Unknown |
| 25 | T68 | $T^{2.39}$ | -4.57 | Unknown |

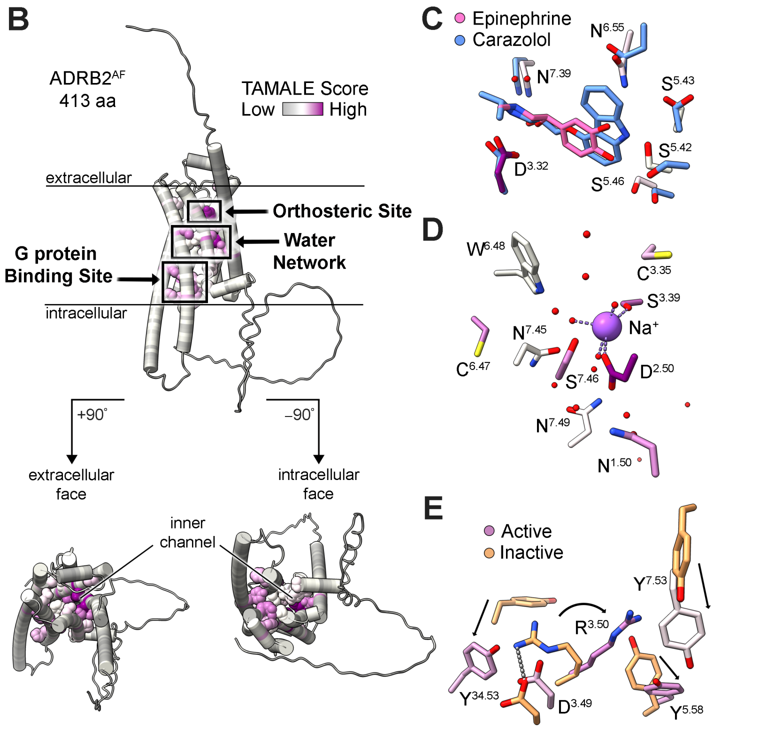


**fig. S13. Prioritization of allosteric features within a GPCR case study.** (**A**) Top 25 TAMALE-predicted ADRB2 functional residues (UniProt ID P07550). (**B**) ADRB2 AlphaFold (AF) model prediction with TAMALE coloring reveals three clusters demarcated by boxes (*1*). (**C**) Overlay of active (TAMALE color) and inactive (blue) states of ADRB2 where the orthosteric site is bound to agonist ligand epinephrine (pink) and antagonist ligand carazolol (blue) (PDB ID 4LDO, 2RH1; (*82, 83*)). (**D**) Sodium-bound site and conserved water network with ADRB2 TAMALE scores superimposed onto consensus allosteric residues of ADRB1 for a proxy due to the absence of a sodium-bound ADRB2 structure (PDB ID 4BVN; (*84*)). (**E**) Active (TAMALE color) and inactive (orange) states of G protein-binding site of ADRB2 (PDB ID 2RH1, 3SN6; (*38, 83*)).

**A** **Las1L (Endoribonuclease)**

| Rank | Residue | Score | Function |
|---|---|---|---|
| 1 | H156 | -0.11 | Active Site |
| 2 | H160 | -0.35 | Active Site |
| 3 | C474 | -1.02 | Unknown |
| 4 | D328 | -1.31 | Unknown |
| 5 | E400 | -1.37 | Unknown |
| 6 | D93 | -1.39 | Unknown |
| 7 | H390 | -1.47 | Unknown |
| 8 | E337 | -1.53 | Unknown |
| 9 | E425 | -1.54 | Unknown |
| 10 | Y266 | -1.66 | Unknown |
| 11 | C97 | -1.96 | Unknown |
| 12 | R155 | -2.02 | Active Site |

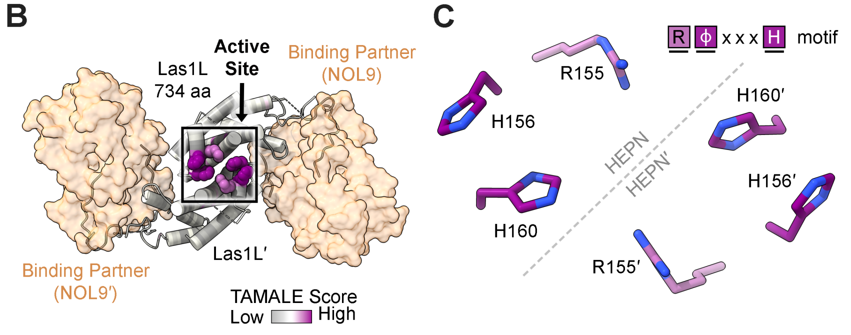


**fig. S14. HEPN catalytic motif prioritization within the Las1L ribonuclease.** (A) Top 12 TAMALE-predicted Las1L functional residues (UniProt ID Q9Y4W2). (**B**) Dimeric Las1L with TAMALE coloring bound to its binding partner (NOL9, orange) reveals a cluster demarcated by a box (*1*). Las1L binds Nol9 for enzyme activation and is shown for reference (*46, 85*). (**C**) Composite Las1L HEPN ribonuclease site formed by two juxtaposed RϕxxxH catalytic motifs (where ϕ is D, N, or H and x is any amino acid) in TAMALE color (PDB ID 9DUN; (*46*)).

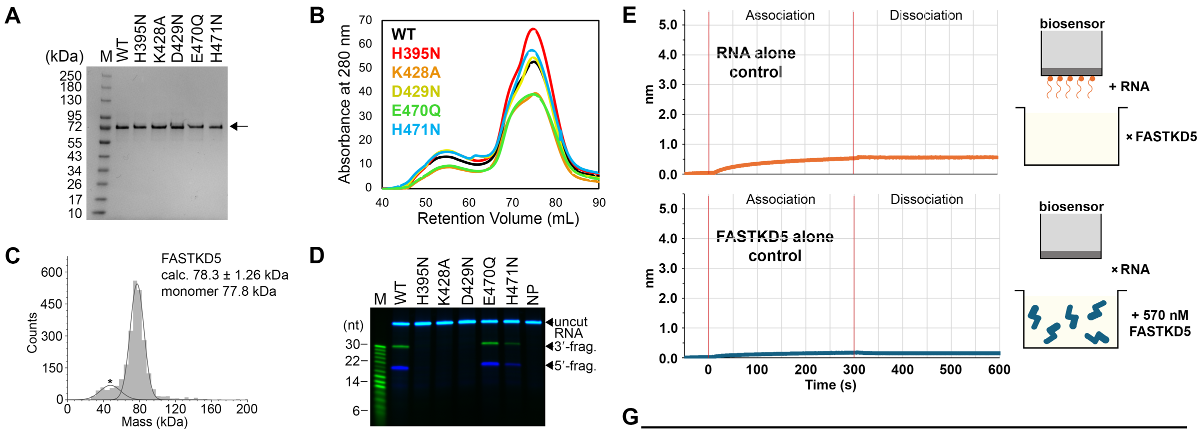


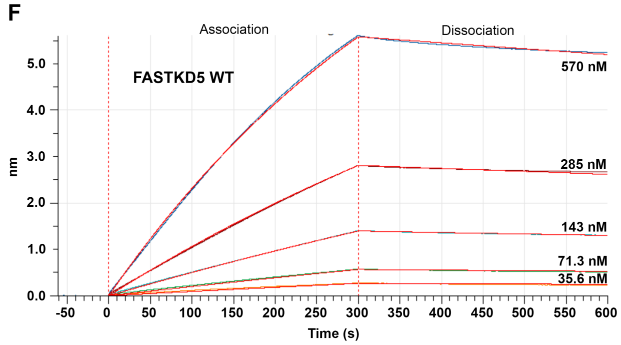


**G**

| FASTKD5 Variant | Rep. i.d. | Kd (nM) | $k_{on}$ ($M^{-1}$ $sec^{-1}$) | $k_{off}$ ($sec^{-1}$) |
|---|---|---|---|---|
| WT | 1 | 63.5 ± 0.260 | $3.7e^{3} \pm 8.4e^{0}$ | $2.4e^{-4} \pm 8.1e^{-7}$ |
| | 2 | 74.3 ± 0.302 | $2.9e^{3} \pm 7.3e^{0}$ | $2.1e^{-4} \pm 7.0e^{-7}$ |
| H395N | 1 | 79.0 ± 0.378 | $2.4e^{3} \pm 7.5e^{0}$ | $1.9e^{-4} \pm 7.0e^{-7}$ |
| | 2 | 62.3 ± 0.372 | $2.7e^{3} \pm 8.7e^{0}$ | $1.7e^{-4} \pm 8.4e^{-7}$ |
| K428A | 1 | 46.0 ± 0.304 | $7.4e^{3} \pm 2.3e^{1}$ | $3.4e^{-4} \pm 2.0e^{-6}$ |
| | 2 | 57.7 ± 0.295 | $5.6e^{3} \pm 1.5e^{1}$ | $3.2e^{-4} \pm 1.4e^{-6}$ |
| D429N | 1 | 75.3 ± 0.358 | $7.0e^{3} \pm 2.2e^{1}$ | $5.3e^{-4} \pm 1.8e^{-6}$ |
| | 2 | 53.7 ± 0.283 | $8.5e^{3} \pm 2.5e^{1}$ | $4.5e^{-4} \pm 2.0e^{-6}$ |
| E470Q | 1 | 65.7 ± 0.346 | $3.5e^{3} \pm 1.1e^{1}$ | $2.3e^{-4} \pm 9.9e^{-7}$ |
| | 2 | 43.3 ± 0.221 | $4.9e^{3} \pm 1.1e^{1}$ | $2.1e^{-4} \pm 9.7e^{-7}$ |
| H471N | 1 | 57.1 ± 0.286 | $1.2e^{4} \pm 3.7e^{1}$ | $6.7e^{-4} \pm 2.6e^{-6}$ |
| | 2 | 49.6 ± 0.230 | $1.3e^{4} \pm 3.3e^{1}$ | $6.3e^{-4} \pm 2.4e^{-6}$ |

**fig. S15. Biochemical characterization of recombinant FASTKD5 variants.** (**A**) SDS-PAGE analysis of purified recombinant FASTKD5 variants along with a broad range protein ladder. (**B**) Size exclusion chromatography profile of recombinant FASTKD5 variants. (**C**) Representative mass photometry analysis of 25 nM wild-type FASTKD5 (total counts 2,019). The molecular mass was determined from three independent measurements to calculate (calc) a mean and standard deviation. The theoretical molecular mass of a FASTKD5 monomer is 77.8 kDa. Asterisk marks the background buffer signal as described previously (*74*). (**D**) Representative urea-PAGE gel of fluorescently labeled ATP8/6-CO3 model RNA (100 nM) cleavage by FASTKD5 variants (50 nM). The RNA substrate is visualized using a 5'-Cy5 fluorophore (blue) and 3'-fluorescein fluorophore (green). A custom fluorescein labeled RNA marker (M) ranging from 30-nucelotides (nt) to 2-nt is shown on the left and a no protein (NP) reaction is a negative control. Three independent replicates were quantified to produce the plot shown in Fig. 5D. (**E**) Representative RNA alone (orange) and FASTKD5 protein alone (blue, 570 nM) control biolayer interferometry sensorgrams. (**F**) Representative biolayer interferometry sensorgram and data fitting of wild-type FASTKD5 (35.6-570 nM) binding activity towards model ATP8/6-CO3 RNA substrate. (**G**) Summary of RNA binding affinity (Kd) and kinetics ($k_{on}$ and $k_{off}$) for FASTKD5 variants. The mean and standard deviation for FASTKD5 binding affinity was computed from two independent replicates and shown in Fig. 5E.

**table S1.**

| | Feature |
|---|---|
| 1 | pos_mean |
| 2 | rest_mean |
| 3 | effect |
| 4 | std |
| 5 | p_value |
| 6 | q_value |
| 7 | sig |
| 8 | sig_p05 |
| 9 | sig_q05 |
| 10 | sig_q05_pos |
| 11 | min_cost |
| 12 | max_cost |
| 13 | min_cost_delta_from_pos_mean |
| 14 | max_cost_delta_from_pos_mean |
| 15 | wcn |
| 16 | rsa |
| 17 | rsa_rel |
| 18 | pae |
| 19 | charge |
| 20 | hydrophobicity |
| 21 | can_act_as_acid |
| 22 | can_act_as_base |
| 23 | x |
| 24 | y |
| 25 | rsa_z |
| 26 | wcn_z |
| 27 | effect_z |
| 28 | seq_n_sig_w5 |
| 29 | seq_frac_sig_w5 |
| 30 | seq_mean_effect_w5 |
| 31 | n_3d_neighbors |
| 32 | n_3d_sig_neighbors |
| 33 | frac_3d_sig_neighbors |
| 34 | mean_3d_effect_neighbors |
| 35 | median_3d_effect_neighbors |
| 36 | max_3d_effect_neighbors |
| 37 | neglog10_p_value |
| 38 | neglog10_q_value |

| | |
|---|---|
| 39 | min_cost_log1p |
| 40 | max_cost_log1p |
| 41 | rsa_log1p |
| 42 | pae_log1p |
| 43 | n_3d_neighbors_log1p |
| 44 | n_3d_sig_neighbors_log1p |
| 45 | seq_n_sig_w5_log1p |
| 46 | Amino acid identity (expanded) |
| 47 | Minimum mutant amino acid pair (expanded) |
| 48 | Maximum mutant amino acid pair (expanded) |

**table S2.**

| Plasmid | Protein Description | Vector | Source |
|---|---|---|---|
| pMP1206 | FASTKD5;<br>HsFASTKD5 wild-type, residues 111-764 | pFastBacHT-B | This study. |
| pMP1219 | FASTKD5 H395N;<br>HsFASTKD5 H395N, residues 111-764 | pFastBacHT-B | This study. |
| pMP1214 | FASTKD5 K428A;<br>HsFASTKD5 K428A, residues 111-764 | pFastBacHT-B | This study. |
| pMP1233 | FASTKD5 D429N;<br>HsFASTKD5 D429N, residues 111-764 | pFastBacHT-B | This study. |
| pMP1225 | FASTKD5 E470Q<br>HsFASTKD5 E470Q, residues 111-764 | pFastBacHT-B | This study. |
| pMP1220 | FASTKD5 H471N;<br>HsFASTKD5 H471N, residues 111-764 | pFastBacHT-B | This study. |

**table S3.**

| Oligo | Substrate | Sequence (5′ to 3′) |
|---|---|---|
| mp2291 | Cleavage substrate | Cy5 - ACCUGCACGACAACACAUAAUGACCCACCAAUCACAUGCCUAUCAUAUAG - Fl |
| mp2329 | BLI substrate | ACCUGCACGACAACACAUAAUGACCCACCAAUCACAUGCCUAUCAUAUAG - Biotin |
| mp1901 | 30-nt marker | Fl - GUUGAGAGAGAGAGAGUUUGAGAGAGAGAG |
| mp2027 | 28-nt marker | Fl - GUUGAGAGAGAGAGAGUUUGAGAGAGAG |
| mp2028 | 26-nt marker | Fl - GUUGAGAGAGAGAGAGUUUGAGAGAG |
| mp2029 | 24-nt marker | Fl - GUUGAGAGAGAGAGAGUUUGAGAG |
| mp2030 | 22-nt marker | Fl - GUUGAGAGAGAGAGAGUUUGAG |
| mp2031 | 20-nt marker | Fl - GUUGAGAGAGAGAGAGUUUG |
| mp2032 | 18-nt marker | Fl - GUUGAGAGAGAGAGAGUU |
| mp2033 | 16-nt marker | Fl - GUUGAGAGAGAGAGAG |
| mp1965 | 14-nt marker | Fl - GUUGAGAGAGAGAG |
| mp1966 | 12-nt marker | Fl - GUUGAGAGAGAG |
| mp1967 | 10-nt marker | Fl - GUUGAGAGAG |
| mp1968 | 8-nt marker | Fl - GUUGAGAG |
| mp1969 | 6-nt marker | Fl - GUUGAG |
| mp1970 | 4-nt marker | Fl - GUUG |
| mp1971 | 2-nt marker | Fl - GU |

Fl defines the fluorescein fluorophore.

**data S1. (separate file)**

This file contains the tabulated UniProt codes for reviewed human enzymes used in the training (train), validation (val), and test sets.
